# Combined lactate- and phosphate-dependent cytoplasmic acidification drives *Mycobacterium tuberculosis* growth arrest at acidic pH

**DOI:** 10.64898/2026.05.15.725484

**Authors:** Adam P. Kibiloski, Shelby J. Dechow, Bassel J. Abdalla, Heather M. Murdoch, Anna D. Tischler, Robert B. Abramovitch

**Affiliations:** Department of Microbiology, Genetics, & Immunology, Michigan State University, East Lansing, MI, 48824; Department of Microbiology and Immunology, University of Minnesota, Minneapolis, MN 55455

## Abstract

*Mycobacterium tuberculosis* (Mtb) cultured in minimal medium at acidic pH arrests its growth when provided specific single carbon sources, including glycerol, propionate, and lactate, a phenomenon we refer to as acid growth arrest. To define mechanisms of acid growth arrest on lactate, transposon mutants that suppress growth arrest were selected. Four mutants had insertions in *phoT* and one had an insertion in *pstC2*, both components of a phosphate ABC transporter. Mtb grows in minimal media supplemented with lactate at acidic pH when phosphate is depleted, showing that Mtb growth arrest on lactate is dependent on phosphate. The combination of lactate and phosphate at acidic pH causes cytoplasmic acidification below pH 6.7 in wild type Mtb, but a *phoT::Tn* mutant maintains a cytoplasmic pH of >7.2. Membrane potential in wild type Mtb is slightly decreased by lactate in a dose-dependent manner but is higher in the *phoT::Tn* mutant. Thus, acidic pH, phosphate, and lactate act together to dissipate proton motive force (PMF), a stress that is associated with acid growth arrest. Transcriptional profiling further supports that lactate causes PMF stress including induction of electron transport chain genes. The *phoT::Tn* mutant grown in lactate at acidic pH upregulates the *senX3*/*regX3* regulon and using a *regX3* mutant, we demonstrate that growth on lactate at low phosphate requires *regX3*. We propose a model where 1) the combined impact of acidic pH, lactate, and phosphate drives cytoplasmic pH acidification and decreased PMF, thus promoting acid growth arrest, and 2) low phosphate or a mutated phosphate transporter causes upregulation of *senX3*-*regX3*, which may induce ESX-5 and PPE/PE-based import mechanisms, thereby altering the mycomembrane or nutrient uptake in a manner that promotes growth on lactate at acidic pH.

**Importance:** *Mycobacterium tuberculosis* (Mtb) grows well on lactate as a sole carbon source at neutral pH, but not at acidic pH. This study sought to understand why there is a pH-dependent growth restriction on lactate. A genetic selection for mutants that can grow on lactate at acidic pH identified mutants defective in phosphate transport. We found that limiting phosphate through depleting extracellular availability or inactivating a phosphate transporter promotes growth on lactate at acidic pH, and that this growth is dependent on the phosphate responsive two-component regulatory system SenX3-RegX3. Furthermore, we show that lactate, phosphate, and acidic pH combine to cause cytoplasmic pH acidification, a metabolic stress that is associated with acid growth arrest on lactate.

## Introduction

*Mycobacterium tuberculosis* (Mtb) is an intracellular bacterial pathogen that survives and replicates in macrophages. Following phagocytosis, Mtb encounters mildly acidic pH in the macrophage phagosome (1). The ability of Mtb to sense and adapt its physiology to acidic pH is necessary for virulence in both the macrophage and animal models (2–7). In defined minimal media supplemented with a single carbon source at acidic pH, Mtb can either grow or establish a state of non-replicating persistence, depending on the carbon source provided (8–11). For example, when Mtb is cultured in minimal medium buffered at pH 5.7 and supplemented with permissive carbon sources that feed the anaplerotic node, such as pyruvate or oxaloacetate, Mtb grows relatively well (8). However, when supplemented with other non-permissive carbon sources at pH 5.7, such as glycerol, lactate, or propionate, Mtb enters a non-replicative state we refer to as acid growth arrest (10).

The acid growth arrest phenotype was initially perplexing, because Mtb has everything necessary to grow under these conditions, including oxygen as a terminal electron acceptor and a carbon source that supports growth at pH 7.0. Therefore, we hypothesized that acid growth arrest is a regulated phenomenon, and that mutants that suppress the acid growth arrest phenotype can be selected. To test this hypothesis, we conducted a genetic selection for mutants that could grow on glycerol as a sole carbon source, and successfully isolated mutants capable of growing on glycerol at acidic pH. We referred to this phenotype as Enhanced Acid Growth (EAG), and this selection indicated that growth restriction on glycerol can be overcome by genetic mutation (12). Notably, all of the isolated mutants with the EAG phenotype were dominant mutations in *ppe51* that increase glycerol uptake to promote enhanced replication both *in vitro* and in activated macrophages (12). Notably, the increased replication in macrophages, resulted in reduced fitness, supporting that integration of acidic pH and carbon source utilization is required to regulate growth and promote pathogenesis in macrophages. We also isolated suppressors of acid growth arrest on propionate as a sole carbon source; all of the mutants were in the *phoPR* two-component system (13). Genetic and biochemical studies show that growth arrest on propionate is due to the diversion of the propionate towards the synthesis of methyl-branched lipids. Thus, mechanisms of acid growth arrest on glycerol and propionate are distinct and point towards carbon-source specific stresses during acid growth arrest. Additionally, *ppe51* EAG variants and *phoPR* mutants could not grow on lactate at acidic pH (12, 13), supporting that yet another mechanism underlies acid growth arrest on lactate.

Mtb catabolizes lactate to promote growth in human macrophages (14), demonstrating its importance during infection. Other pathogens such as *Neisseria sp*., *Hemophilus influenzae*, and *Staphylococcus aureus* also use host-derived lactate to promote pathogenesis (15). In this study, we aim to define the mechanism of acid growth arrest of Mtb cultured at acidic pH with lactate as the sole carbon source. Towards this goal, we conducted a genetic selection for suppressor transposon mutants that grow on lactate at acidic pH. The selection successfully identified several mutants, including four mutants in *phoT* and one mutant in *pstC2*. Both PhoT and PstC2 are components of phosphate ABC transporter system PstSCA-PhoT, supporting a link between phosphate and growth arrest on lactate. We found that growth arrest on lactate is dependent on phosphate concentrations and is also dependent on the two-component regulatory system SenX3-RegX3. We also found that lactate and phosphate negatively impact PMF through cytoplasmic acidification. Together, findings from Tischler *et al*. (16), Healy *et al*., (17) and this study, reinforce the critical role for phosphate in regulating pH-dependent adaptations.

## Results

### Mtb is nonreplicating and viable on lactate at acidic pH

Previous studies found that wild type (WT) Mtb is unable to grow on lactate at pH 5.7 (8, 12). In defined medium with lactate as the sole carbon source WT Erdman grows at pH 7.0, but arrests growth at pH 5.7 (Fig. 1A). We hypothesized the arrested bacteria are viable at pH 5.7, but not replicating. To test this hypothesis, following 9 days of growth arrest on 4 mM lactate, the permissive carbon source pyruvate was added to the culture. At day 14 the bacteria began to replicate again, showing they remain viable on lactate at pH 5.7 (Fig. 1A). To confirm viability, we enumerated Mtb CFUs, and observed no reduction in CFUs over time. (Fig. 1B). This pattern of growth was also observed in WT CDC1551 (Fig. S1A). Therefore, acid growth arrest on lactate is a state of non-replicating persistence.

**Figure 1.**
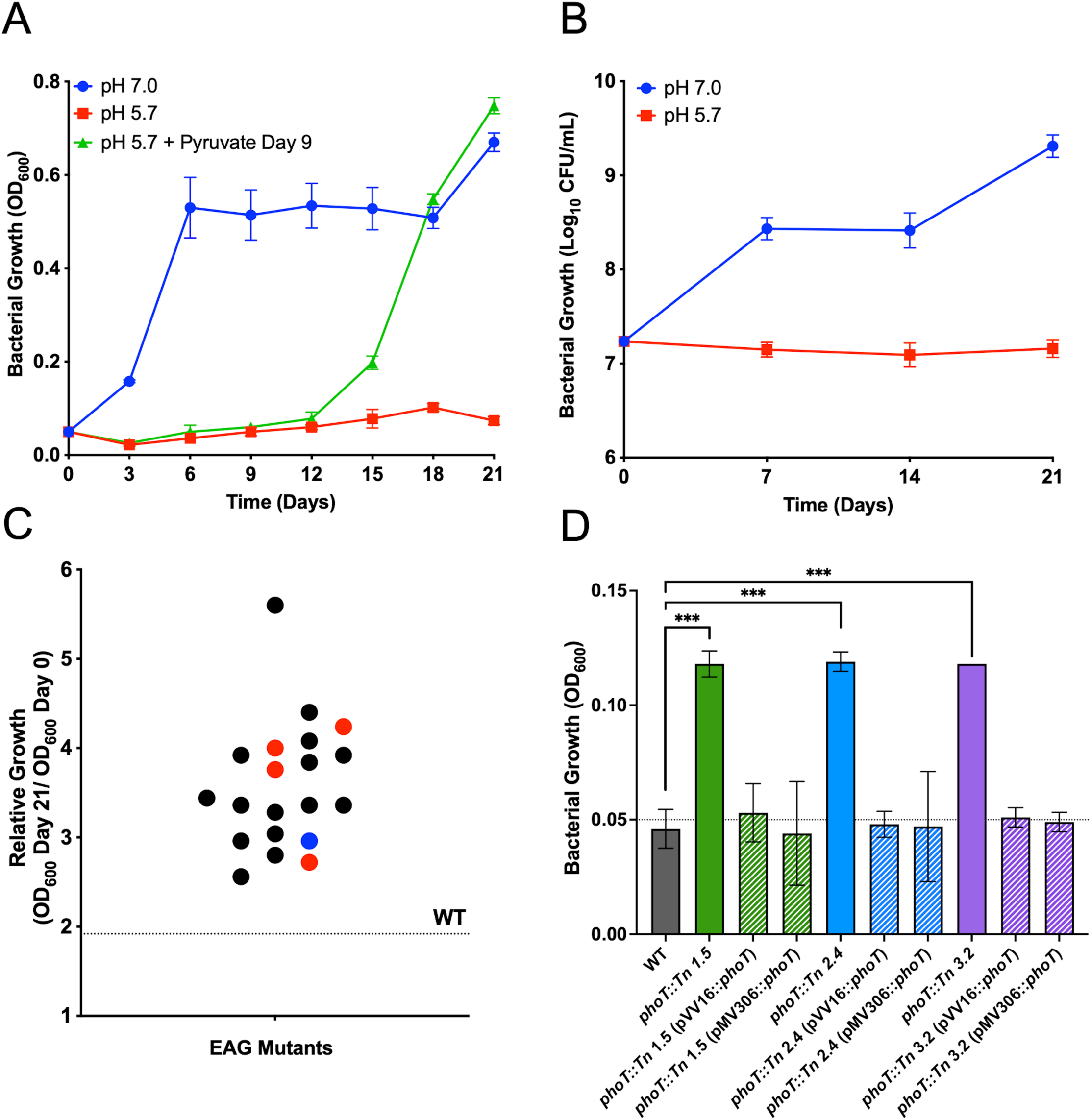
Selection of *phoT* mutants that suppress acid growth arrest on lactate. A) WT Erdman grown in minimal medium buffered to pH 5.7 (red) and 7.0 (blue) supplemented with 4 mM lactate. On day 9, 10 mM pyruvate was added to Mtb grown on lactate at pH 5.7 (green). This experiment was performed at least twice with similar results. Error bars indicate standard deviation of three technical replicates. B) WT Erdman grown in minimal medium buffered to pH 5.7 (red) and pH 7.0 (blue), supplemented with 4 mM lactate. This experiment was performed at least twice with similar results. Error bars indicate the standard deviation of three technical replicates. C) Confirmation of suppressor mutants by examining relative growth at Day 21 OD_600_ to OD_600_ from the initial Day 0 inoculum. Each dot represents a selected transposon mutant isolated from MMAT agar plates and confirmed for enhanced growth in liquid MMAT (pH 5.7) supplemented with 4 mM lactate. Red dots indicate *phoT::Tn* mutants, the blue dot represents a *pstC2::Tn* mutant. The dotted line indicates the relative WT Erdman growth (> ratio of 1). This experiment was performed once in single replicate. D) Three of the *phoT::Tn* mutants were complemented using a single copy chromosomal insertion vector (pMV306) or an episomal overexpression vector (pVV16). Black bar indicates growth arrest in WT. Data is OD_600_ after 21 days of growth. Statistical analysis was performed using an unpaired t-test (***, p<0.001). This experiment was performed once. Error bars indicate the standard deviation of three technical replicates.

### Selection of mutants that suppress growth arrest on lactate at pH 5.7

In minimal medium at acidic pH with lactate, Mtb has everything it needs to grow, including a carbon source and oxygen. Therefore, we hypothesized that acid growth arrest on lactate is a regulated adaptation and that suppressor mutants could be selected. To test this hypothesis, we conducted a selection for suppressor mutants by plating WT Mtb, at a density of 10^9^ CFUs on plates buffered to pH 5.7 and supplemented with 4 mM lactate as a sole carbon source. Despite >8 weeks of incubation at 37°C, no spontaneous mutants were isolated. Notably, this is the same method we used to generate *ppe51* variants on glycerol, which appeared at a high frequency of resistance of 10^-6^ to 10^-7^ (12). To increase the number of mutants examined, we repeated the selection using a Mtb Erdman transposon mutant library containing >100,000 mutants, at a density of 10^9^ CFUs per plate. After eight weeks of incubation, we observed >1000 colonies across four plates, of which 20 colonies were randomly selected for further confirmation. All 20 mutants exhibited increased growth on lactate at pH 5.7 compared to WT Erdman, which was growth arrested (Fig. 1C and S1B). The transposon insertion sites of 15 mutants were identified using either inverse PCR (18) and Sanger sequencing or whole genome sequencing (Table 1). We identified four independent transposon insertions in *phoT* (*rv0820*), which encodes an ATP-binding protein phosphate ABC transporter, and one transposon mutant with an insertion in *pstC2* (*rv0929*), which encodes a phosphate ABC transporter integral membrane protein. Additionally, we observed single transposon insertions in *fadD17*, *lppF*, *rv1056*, *fadD4*, and *modB*. Single intergenic insertion sites were also observed between *rv3226c* and *aroA*; *cyp136* and *rv3060c*; and *rv2929* and *fadD26* (Table 1). *phoT::Tn* strains were complemented by cloning WT *phoT* in either a single-copy chromosomal insertion vector (pMV306::*phoT*) or an episomal overexpression vector (pVV16::*phoT*). When *phoT::Tn* mutant strains 1.5, 2.4, or 3.2 and their respective complemented strains were grown in minimal medium at acidic pH amended with 4 mM lactate, the WT and all complemented strains exhibited growth arrest while the transposon mutants exhibited growth (Fig. 1D). These results confirm that the loss-of-function transposon insertions in *phoT* promote growth on lactate. Going forward, the *phoT::Tn* strain 1.5 was used for the remainder of the studies unless otherwise specified.

**Table 1.** Transposon insertion sites that permit growth on lactate. Mutants were arbitrarily assigned identification numbers based on the order they were picked from the plate. FC indicates fold-change increased growth compared to WT on lactate. TA site indicates the TA in which the transposon recognized and inserted. The numbers indicate the nucleotide number the TA is at over the total nucleotide length of the gene. The method of identification is either whole-genome sequencing (WGS) or inverse PCR as indicated in column five. Five mutants had no identifiable transposon insertion.

| Mutant # | Insertion Start Site | Insertion Stop Site | Approx TA site/gene length (nt) | FC relative to WT growth | Rv Number | Gene Name | Method of identification |
| --- | --- | --- | --- | --- | --- | --- | --- |
| 1.1 | 3937415 | 3937416 | TA 584/1509 | 3.04 | Rv3506 | *fadD17* | WGS |
| 1.2 | N/A | N/A | No transposon identified | 3.92 | N/A | N/A | WGS |
| 1.3 | 2769395 | 2769398 | intergenic | 3.36 | N/A | N/A | WGS |
| 1.4 | 2173253 | 2173254 | TA - 1/1272 | 2.56 | Rv1921c | *lppF* | WGS |
| 1.5 | 916505 | 916506 | TA 197/777 | 2.72 | Rv0820 | *phoT* | Inverse PCR |
| 2.1 | 1829378 | 1829379 | TA 745/2097 | 2.80 | Rv1633 | *uvrB* | Inverse PCR |
| 2.2 | 1183021 | 1183022 | TA 575/765 | 5.60 | Rv1056 | *Rv1056* | WGS |
| 2.3 | 256650 | 256651 | TA 1178/1518 | 3.28 | Rv0214 | *fadD4* | WGS |
| 2.4 | 917077 | 917078 | TA 770/777 | 3.76 | Rv0820 | *phoT* | WGS |
| 2.5 | N/A | N/A | No transposon identified | 2.96 | N/A | N/A | WGS |
| 3.1 | 2104922 | 2105025 | 79/795 | 3.36 | Rv1858 | *modB* | WGS |
| 3.2 | 917060 | 917061 | TA 754/777 | 4.24 | Rv0820 | *phoT* | Inverse PCR/WGS |
| 3.3 | 3608788 | 3608791 | 729/ 759 | 4.40 | Rv3226c | Rv3226c | WGS |
| 3.4 | N/A | N/A | No transposon identified | 3.92 | N/A | N/A | WGS |
| 3.5 | 3427375 | 3427378 | intergenic | 3.36 | N/A | N/A | WGS |
| 4.1 | 3251680 | 3251783 | intergenic | 4.08 | N/A | N/A | Inverse PCR/WGS |
| 4.2 | 917010 | 917013 | 703/777 | 4.00 | Rv0820 | *phoT* | WGS |
| 4.3 | N/A | N/A | No transposon identified | 3.84 | N/A | N/A | WGS |
| 4.4 | N/A | N/A | No transposon identified | 3.44 | N/A | N/A | WGS |
| 4.5 | 1040405 | 1040406 | TA 690/975 | 2.96 | Rv0929 | *pstC2* | WGS |

#### The *phoT::Tn* mutant grows on lactate at acidic pH

We previously observed that the *ppe51* variants and *phoPR* mutants have selective enhanced growth on glycerol or propionate, respectively, and did not permit growth on other non-permissive carbon sources, including lactate (12, 13). We hypothesized that the *phoT::Tn* mutant would also have carbon source specific enhanced growth at acidic pH. To test this hypothesis the WT, *phoT::Tn* and the complemented strains were grown in minimal medium at acidic pH supplemented with individual carbon sources. The only carbon source that the *phoT::Tn* mutant could grow on at acidic pH was lactate (Fig. 2A) with an ∼8-fold increase in growth of the *phoT::Tn* mutant compared to WT. These findings demonstrate that lactate-specific metabolic properties, as compared to the other non-permissive carbon sources, drives growth arrest at acidic pH.

**Figure 2.**
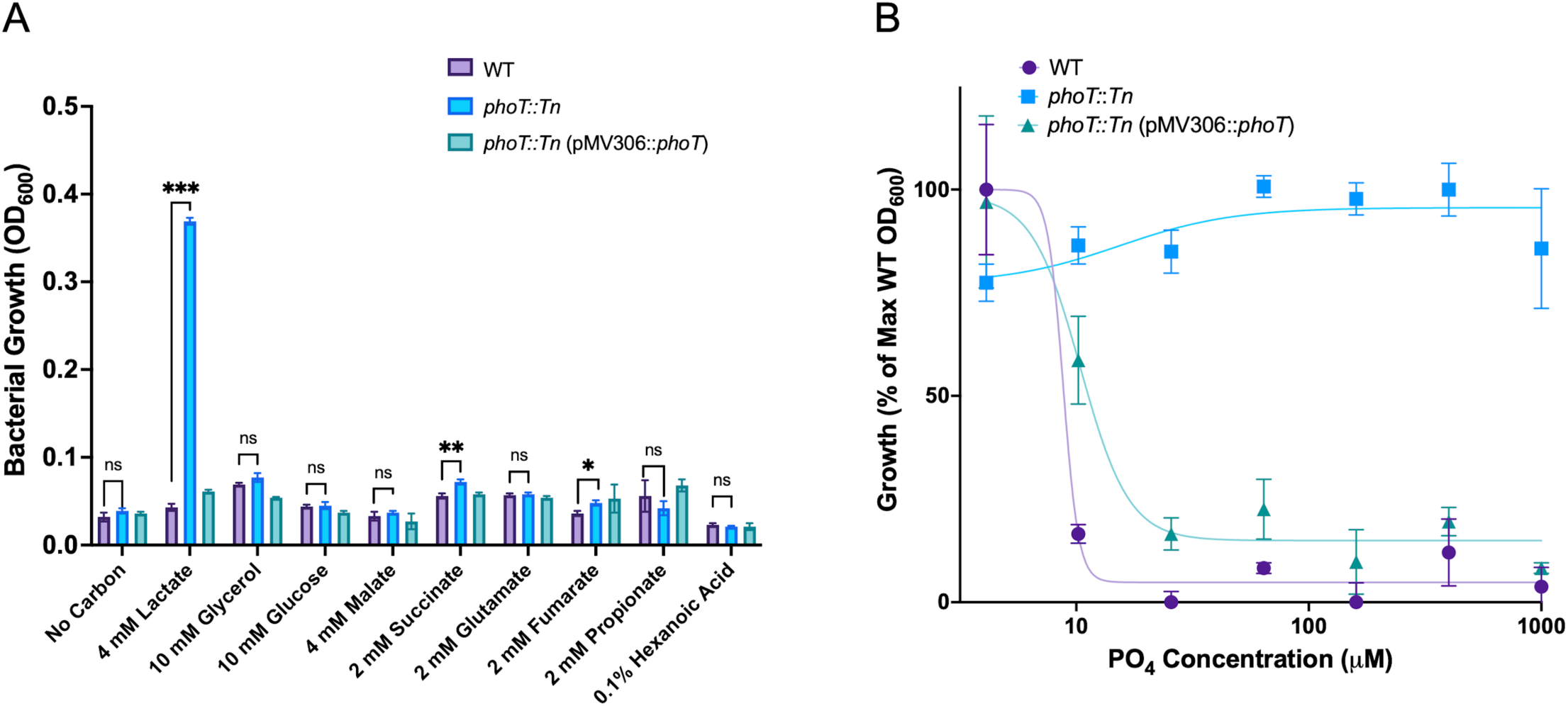
Growth of the *phoT::Tn* mutant is specific to lactate and independent of phosphate concentration. A) Growth of the WT, *phoT::Tn* mutant and the pVV16 complemented strains cultured in a variety of host associated carbon sources. Data measured on day 21 with no growth indicated by the “No Carbon” control. Statistical analysis was performed using an unpaired t-test (**, p<0.005, ***,p<0.001). This experiment was performed at least twice with similar results. Error bars indicate the standard deviation of three technical replicates. B) WT (Purple), *phoT::Tn* (blue), and *phoT::Tn* (pMV306*::phoT*) (teal) strains were grown in MMAT with lactate as the sole carbon source at pH 5.7 across a 2.5-fold dose response of KH_2_PO_4_. Values were normalized to percentage of maximum WT growth. This experiment was performed twice with similar results. Error bars indicate the standard deviation of three technical replicates.

### Phosphate limitation restores Mtb growth on lactate at pH 5.7

We hypothesized that the *phoT::Tn* mutant may have reduced phosphate uptake and that low phosphate may promote growth on lactate at acidic pH. To test this hypothesis, we examined the growth of the WT, *phoT::Tn* mutant and complemented strains grown at pH 5.7 and supplemented with 4 mM lactate across a serial dilution of KH_2_PO_4_ from 4.1 µM to 1.0 mM, along with a no phosphate control. The standard concentration of phosphate in MMAT minimal medium is ∼25 mM. The *phoT::Tn* mutant was insensitive to phosphate, growing well at all tested phosphate concentrations (Fig. 2B). The WT and complemented strains arrested growth at phosphate concentrations at or above 10 µM. However, at phosphate concentrations at 4.1 µM and in the no phosphate control, the WT and complemented strains grew as well as the *phoT::Tn* strain (Fig. 2B). Similarly, in the H37Rv strain, low phosphate promotes growth on lactate at acidic pH (Fig. S2). Therefore, phosphate restricts growth on lactate at acidic pH, but limiting phosphate uptake by disrupting *phoT* or depleting the media of phosphate overcomes the constraint on growth on lactate.

We hypothesized that phosphate and lactate may interact to regulate growth at acidic pH and sought to define the relative concentrations at which growth is permitted or restricted. To define these interactions we conducted checkerboard assays, where Mtb growth was examined across a 2.5-fold serial dilution of lactate ranging between 164 µM – 40 mM with phosphate concentrations ranging from 0.8 µM – 200 µM and a no phosphate control. The only lactate concentrations that permitted growth of any strains in this experiment were in the range between 1 mM and 6.4 mM (Fig. 3A-3D), suggesting a narrow range of permissive lactate concentrations. Lactate concentrations above 6.4 mM were likely unable to drive growth due to associated toxicity (19). Lactate concentrations at and below 1.0 mM were not sufficient to support growth. At the threshold of >12 µM phosphate, only the *phoT* mutant could grow on lactate (Fig. 3A-D, S3A-D, S4). However, at <12 µM phosphate, WT Mtb could grow at 6.4 mM lactate. Thus, it is the combined interplay of phosphate and lactate concentration that drives the acid growth arrest phenotype. Additionally, pH dependence was determined by growing WT, *phoT::Tn*, and pMV306-complemented strains in a range of pHs between 5.5 and 7.0 on 4mM lactate. The *phoT::Tn* mutant exhibited growth at pH 5.5-6.0, with the most robust growth observed at pH 6.0, growing 10-fold better than the WT (Fig. S5A). At pH 6.5 or greater, robust growth was observed for all the strains. These data together demonstrate that lactate concentration, phosphate concentration, and acidity contribute to growth regulation.

**Figure 3.**
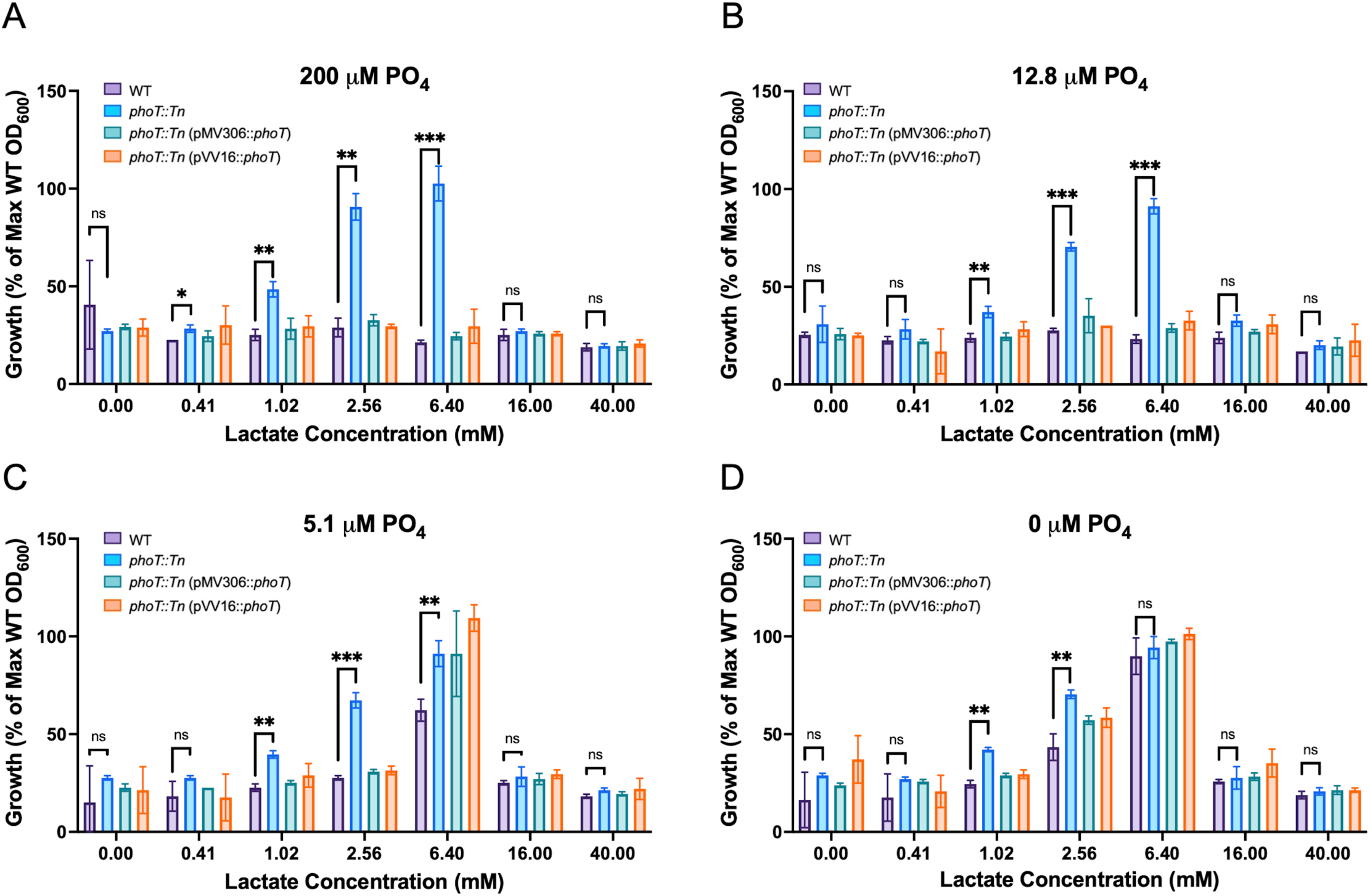
Mtb growth arrest on lactate is dependent on phosphate concentration. Mtb grown at varying lactate concentrations in media amended with (A) 200 µM, (B) 12.8 µM, (C) 5.1 µM, and (D) with no PO_4_. Data is reported as % of maximum of WT growth (observed in the 0 µM phosphate condition). Statistical analyses were performed using an unpaired t-test (*, p<0.05 **, p<0.005 ***, p<0.001) This experiment was performed twice with similar results. Error bars represent the standard deviation of three replicates.

#### Lactate and phosphate function together to cause cytoplasmic acidification

Even under acidic conditions, Mtb maintains a relatively neutral cytoplasmic pH of ∼7.2-7.4 (5, 8). Loss of cytoplasmic pH homeostasis is associated with cell death, possibly due to disruption of the PMF or protein unfolding (5, 20). Lactic acid bacteria exposed to high concentrations of lactate experience a range of stresses including an acidified cytoplasm and changes in oxidative phosphorylation due to PMF stress (21, 22). Both lactate and phosphate are ions that could carry a proton into the cytoplasm when imported for metabolism. This could acidify the cytoplasm or increase membrane potential if the protons are pumped out of the cell to maintain pH-homeostasis. Therefore, we hypothesized that Mtb may be experiencing an acidified cytoplasm or higher membrane potential when exposed to lactate or phosphate. Cytoplasmic pH was examined in Mtb incubated at pH 5.7 in minimal medium with lactate at 0, 1, 4 or 10 mM and at either 0 or 200 µM phosphate. These conditions cover growth arresting or permissive conditions as determined in the studies presented in Figure 3. In the absence of phosphate, lactate causes acidification of the cytoplasm in a dose-dependent manner (Fig. 4). The pH drops from ∼7.2 to ∼6.8 when the bacteria are incubated in 0, 1 or 4 mM lactate with no added phosphate (Fig. 4A-C). Notably, these are growth permissive conditions (Fig. 2B). At 10 mM lactate and 0 µM phosphate, we observe a more acidified cytoplasm of ∼6.4 (Fig. 4D) which may be associated with growth restriction that we observe at high lactate concentrations (Fig. 3). Therefore, lactate causes cytoplasmic acidification in a dose-dependent manner. When 200 µM phosphate is added, we observe a relatively neutral pH at 0 and 1 mM lactate (Fig. 4A-B). However, at 4 mM lactate and 200 µM phosphate, cytoplasmic pH is more strongly acidified below pH 6.5 (Fig. 4C, S5C). These conditions are associated with growth arrest of Mtb (Fig. 2B). The cytoplasmic acidification is dependent on lactate, as we observed a cytoplasmic pH of ∼7.4 in Mtb treated in glycerol or pyruvate as previously published (8) and repeated here (Fig. S5B). Thus, cytoplasmic acidification below a threshold of ∼pH 6.8 is associated with growth arrest, and the acidification requires the combined action of phosphate and lactate.

**Figure 4.**
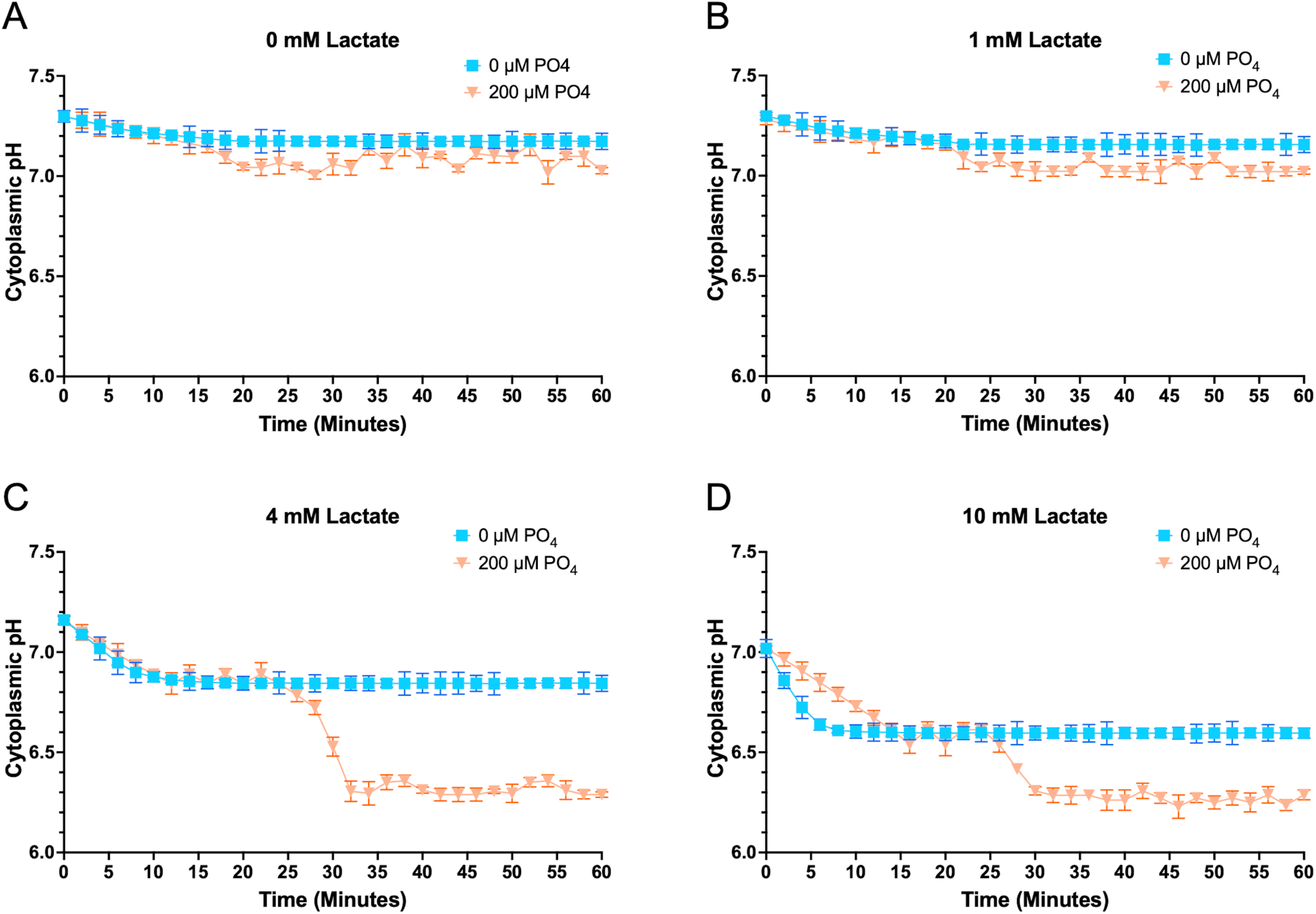
Combined lactate and phosphate cause cytoplasmic acidification. Cytoplasmic pH was observed over time when exposed to 200 µM phosphate (orange), or no phosphate (light blue) in medium at pH 5.7 with no lactate (A), 1 mM lactate (B), 4 mM lactate, and 10 mM lactate (D). This experiment was performed at least twice with similar results. Error bars represent the standard deviation of three technical replicates.

We further hypothesized that because phosphate contributes to cytoplasmic acidification, a *phoT*::Tn mutant may have a more neutral cytoplasm due to lower phosphate import. Using similar concentrations of lactate and phosphate as in the previous experiment, we examined cytoplasmic pH of the WT, *phoT::Tn*, and *phoT::Tn* (pMV306:*:phoT*) strains. In the WT, we saw a similar trend of acidification at higher phosphate and lactate concentrations (Fig 5A). Supporting our hypothesis, the *phoT::Tn* mutant cytoplasm did not decrease below pH ∼7.0 (Fig. 5B). The complemented strain replicated WT cytoplasmic pH trends, with the exception of the 4 mM lactate and no added phosphate condition, where we observed acidification in the complemented strain (Fig 5C). Averages of endpoint data at pH 5.7 (Fig. 5D) support the conclusion that the transposon insertion in *phoT* significantly reduces cytoplasmic acidification. These data reinforce our hypothesis that a cytoplasmic pH below pH 6.8 is associated with growth arrest. This experiment was also conducted in Mtb cultured at pH 7.0, with no cytoplasmic acidification observed below pH 7.2 (Fig 5E, Fig. S6). These data demonstrate that 1) at acidic pH, phosphate, and lactate function together to cause cytoplasmic acidification, and that 2) mutating *phoT* is sufficient to alleviate this stress and suppress acid growth arrest on lactate.

**Figure 5.**
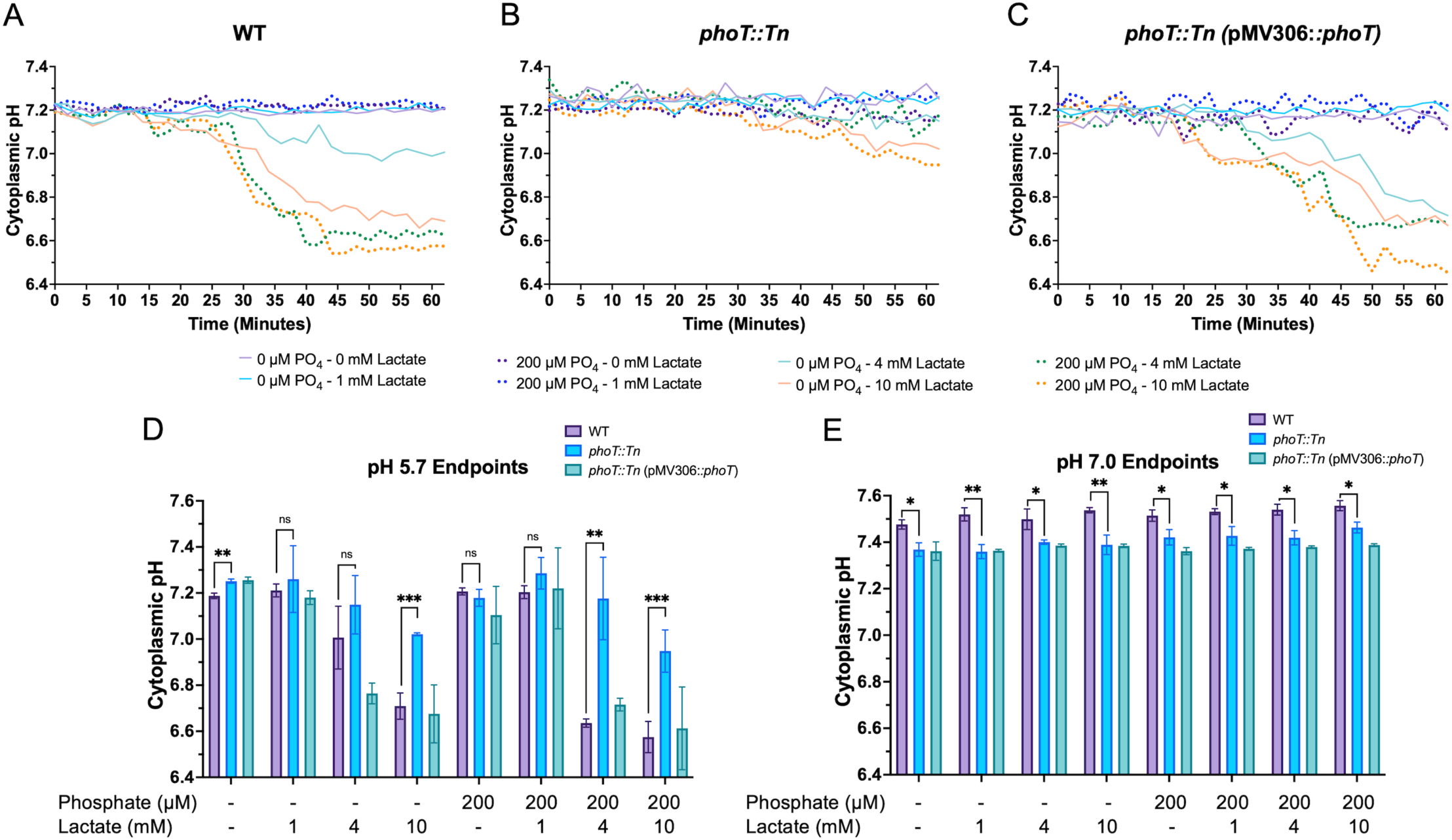
*phoT* mutation alleviates cytoplasmic acidification. Cytoplasmic pH was measured over time in medium with the indicated range of phosphate and lactate concentrations for WT (A), *phoT*::*Tn* (B) and *phoT*::*Tn* (pMV306::*phoT*) (C) strains. Endpoint cytoplasmic pH (60 min) of strains exposed to lactate at pH 5.7 (D) or pH 7.0 (E) Endpoint data of strains exposed to lactate at pH 7.0. Statistical significance determined by an unpaired t-test (**, p<0.005, ***, p<0.001) All experiments for this figure were performed at least twice with similar results. Error bars represent standard deviation of three technical replicates.

#### Lactate and disruption of *phoT* increase membrane potential

Significant cytoplasmic acidification is observed with 200 µM phosphate at 4 mM lactate but not 1 mM lactate. This suggests that Mtb can maintain pH-homeostasis at 1 mM lactate, possibly by the export of protons. This activity is hypothesized to lead to a higher membrane potential. Similarly, restricting phosphate uptake in the *phoT* mutant is also predicted to increase membrane potential. Using similar combinations of lactate and phosphate concentrations as in the cytoplasmic pH assay, we measured membrane potential over time at pH 5.7. Lactate at 1 mM and 4 mM caused a modest increase in membrane potential, consistent with our hypothesis that maintenance of cytoplasmic pH homeostasis would increase membrane potential (Fig. 6A-B, S7A-B). Unexpectedly, high or low phosphate did not impact membrane potential, at acidic or neutral pH (Fig 6A-B and S8A-S8B), suggesting that its primary impact is on cytoplasmic acidification. At lower phosphate levels (0-200 µM), we observed minor changes in membrane potential in the *phoT::Tn* mutant, at acidic and neutral pH (Fig. S7A-B). From these data, we conclude that the *phoT::Tn* mutation does not have a substantial impact membrane potential at the tested concentrations of lactate and phosphate.

**Figure 6.**
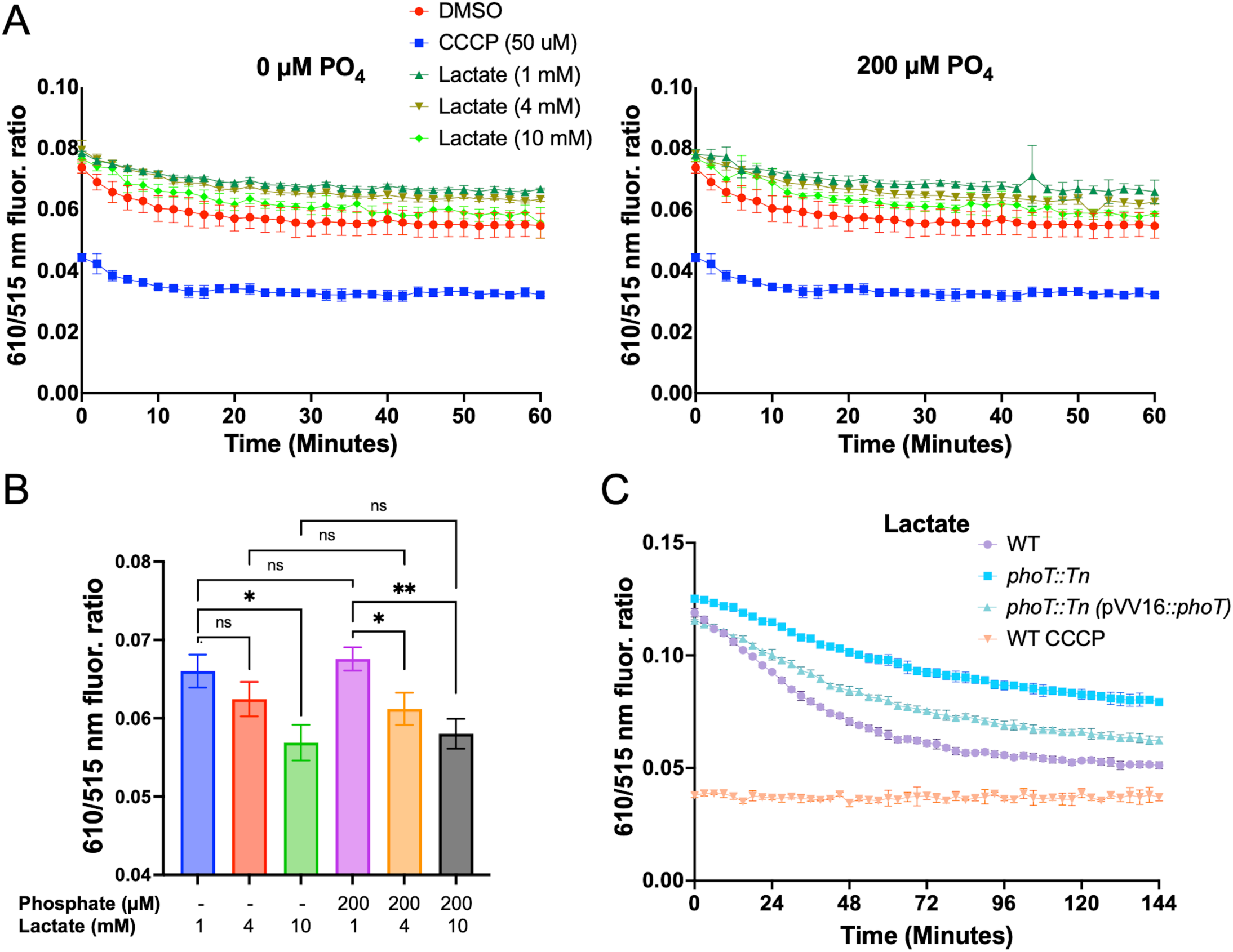
Lactate decreases membrane potential and the *phoT::Tn* has higher membrane potential. A) Membrane potential was evaluated over time at 0 µM phosphate, and 200 µM phosphate. Lactate concentrations tested were 10 mM (light green), 4 mM (yellow-green), and 1 mM (forest green). A DMSO vehicle control was included (red). This experiment was performed twice with similar results. Error bars indicate the standard deviation of three technical replicates. B) Minutes 46-60 were averaged from panel A. These data represent endpoint data from the same experiment. C) Membrane potential of strains incubated in MMAT medium with different carbon sources. The strains were evaluated were *phoT::Tn* (pVV16::*phoT*) (teal), *phoT::Tn* (light blue), and WT (purple). WT treated with CCCP (orange) was used as a depolarized control. This experiment was performed at least twice with similar results. The error bars represent the standard deviation of three technical replicates.

We investigated membrane potential of WT, *phoT*::*Tn*, and complemented strains across different carbon sources in MMAT medium, which has a high concentration of phosphate (25 mM). In the *phoT::Tn* mutant on lactate, membrane potential was substantially higher than the WT or complemented strains (Fig. 6C), demonstrating that exclusion of phosphate can lead to a higher membrane potential at high phosphate. The higher membrane potential of *phoT::Tn* mutant was also observed in MMAT with either pyruvate or glycerol at pH 5.7 (Fig. S8). We conclude that at lower phosphate concentrations (up to 200 µM), the primary impact of lactate and phosphate is on cytoplasmic acidification. However, at high concentrations of phosphate (25 mM in unmodified MMAT), blocking phosphate uptake leads to an increase in membrane potential. Given that under the conditions examined in Figures 4, 5, and 6A, growth arrest is associated with a lower cytoplasmic pH and mostly unchanged membrane potential, we conclude that growth arrest is associated, and potentially caused by, decreased PMF.

#### The *phoT::Tn* mutant is sensitive to ETC targeting agents on lactate with high phosphate

Given the impact of lactate and phosphate on PMF, we hypothesized that growth arrest may be caused by dysregulation of pH-homeostasis and oxidative phosphorylation functions of the electron transport chain (ETC). To test this hypothesis, we examined the sensitivity of Mtb to known ETC-targeting compounds, under different carbon sources (pyruvate or lactate), pHs (pH 7.0 or 6.0), and phosphate concentrations (0 and 200 µM phosphate). We used pH 6.0 instead of pH 5.7 to improve the dynamic range of the growth in the 96-well plate format. ETC targeting compounds were tested across a dose response and include, thioridazine (TDZ) that targets the non-proton pumping NDH-2 (23), clofazimine (CLZ) that targets NDH-2 to generate reactive oxygen species (ROS) among other mechanisms (24–26), Q203 which targets cytochrome *bcc::aa_3_* complex (27, 28), bedaquiline (BDQ) that targets ATP synthase (29–31), nigericin (NIG) that acts as a K^+^/H^+^ antiporter that disrupts the proton gradient (32–34), CCCP that collapses PMF by shuttling protons into the cell (35–37), and valinomycin (VAL) that binds and carries K^+^ ions out of the cell and collapses membrane potential (38–40).

Mtb exhibited differential sensitivity to several ETC inhibitors depending on the growth conditions and *phoT* mutation, supporting impacts of phosphate, lactate and pH on modulating the ETC (Fig. S9). The *phoT::*Tn mutant was more sensitive to all ETC targeting agents in lactate at high phosphate, as compared to the WT or complemented strains. Given that the sensitivity of the *phoT*::*Tn* mutant is not pH-dependent, and that Mtb in pyruvate are more resistant, the sensitivity is likely not simply driven by enhanced growth. Rather, the fact that the enhanced sensitivity is lost on lactate at low phosphate, this suggests that the combined action of lactate and high phosphate in the *phoT::Tn* mutant is selectively causing changes to the ETC or PMF. In the WT, we also observed specific sensitivities to ETC inhibitors, consistent with combined impacts of lactate and phosphate on ETC function. For example, Mtb exhibited phosphate and lactate dependent resistance to valinomycin, with an almost complete loss of activity at lactate at pH 6.0 or 7.0 in the presence of 200 µM phosphate. In comparison, growth on pyruvate with phosphate, or lactate without phosphate, promoted valinomycin sensitivity. Several drugs demonstrated enhanced activity in lactate at pH 6.0 as compared to pH 7.0, with the effect independent of phosphate, including clofazimine, nigericin, Q203 and CCCP, consistent with acidic pH sensitizing Mtb to ETC targeting drugs. Overall, these data show that the *phoT::Tn* has enhanced sensitivity to ETC targeting agents selectively on lactate in high phosphate conditions, supporting dysregulation of PMF by the combined action of lactate and phosphate.

### Transcriptional profiling defines adaptations associated with lactate-dependent growth arrest

Both lactate and propionate provided as a single carbon source arrest growth at acidic pH. In a recent study, we found that a *phoPR* knockout mutant can grow on propionate at acidic pH (13). These observations enable the comparison of transcriptional profiles of acid growth arrest on two different non-permissive carbon sources. In the propionate study, we generated a *phoR::Tn* mutant from a selection on propionate at acidic pH. To understand how the WT, *phoT::Tn* and *phoR::Tn* mutant strains respond to acidic pH in the presence of lactate or propionate, RNA-sequencing transcriptional profiling was conducted. WT Erdman, the *phoT::Tn,* and *phoR::Tn* mutant were cultured in MMAT buffered to pH 7.0 or pH 5.7 supplemented with 4 mM lactate or 2 mM propionate for 3 days, after which RNA was extracted, and global transcriptional profiles were analyzed (Tables S1-S5).

Comparisons of WT Mtb cultured on lactate versus propionate at acidic pH, shows shared responses to acidic pH and distinct responses to specific carbon sources. For example, when examining differentially expressed genes on lactate or propionate at pH 5.7 vs pH 7.0, 202 and 205 of the upregulated and downregulated genes, respectively, were shared between the two carbon sources (Fig 7C). However, greater than 200 genes were significantly and differentially upregulated in each respective carbon source (Fig. 7C). That is, growth arrest on propionate is distinct from growth arrest on lactate. The overlap between the transcriptional signatures of these two carbon sources is likely associated with adaptation to acidic pH, including PhoPR regulated genes (Fig. 7A). Genes that are only upregulated in lactate at acidic pH include those associated with the electron transport chain, metals uptake, phosphate import, and arginine biosynthesis. Particularly a large number of cytochromes were upregulated including *cydBCD* (Fig. 7A).

**Figure 7.**
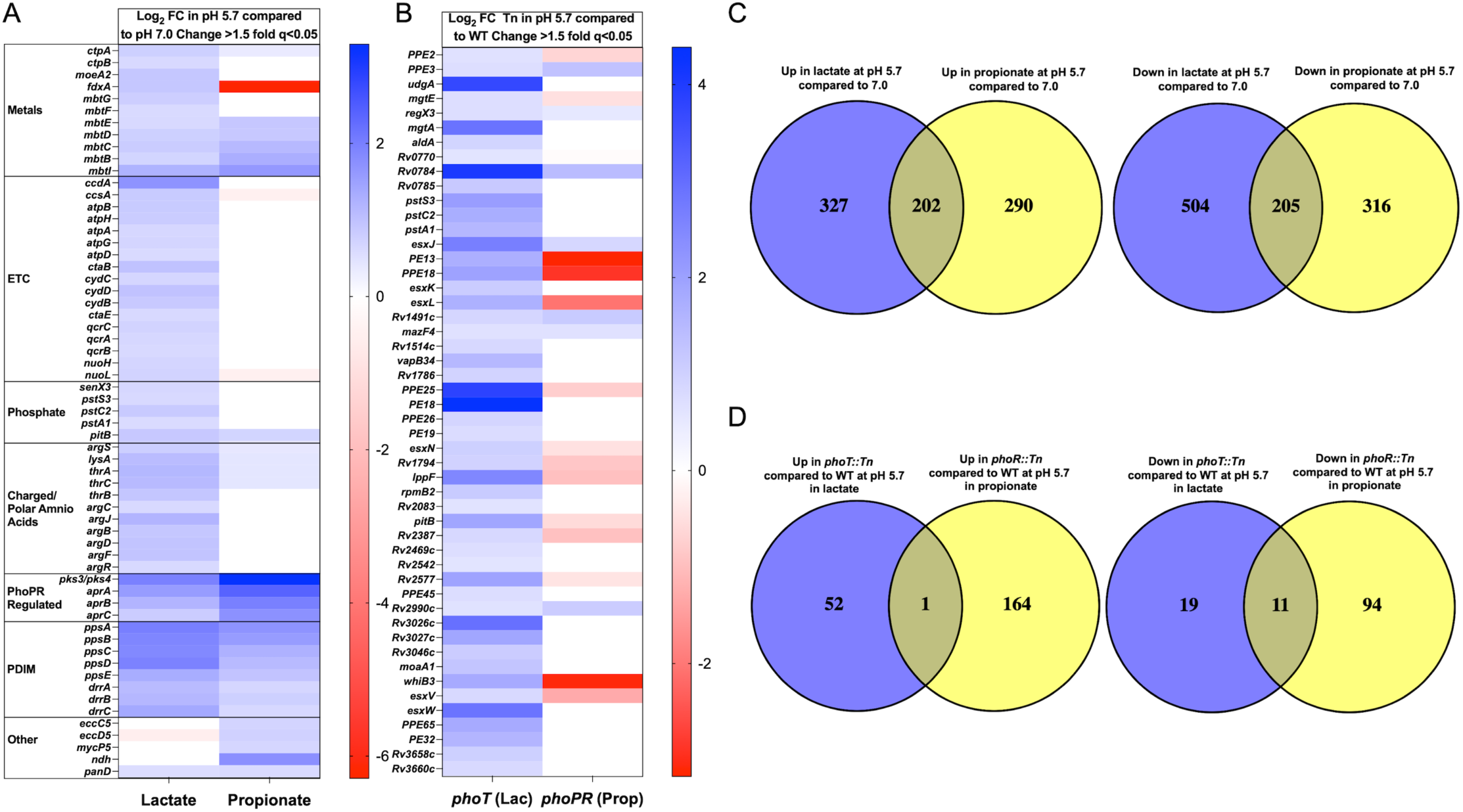
Differential gene expression in lactate vs. propionate. A) Heat map of genes differentially expressed (fold change >1.5x and q<0.05) in WT when exposed to different carbon sources. Genes up and downregulated in lactate at pH 5.7 compared to pH 7.0 on the left. Genes up and downregulated in propionate at pH 5.7 compared to pH 7.0 on the right. B) Heat map of genes differentially expressed (fold change >1.5x and q<0.05) in mutant strains capable of growing on their carbon sources at pH 5.7 compared to WT in the same carbon source as each respective mutant at pH 5.7. Genes that change their expression in this comparison indicate how growth affects gene expression compared to growth arrest. C) Venn diagrams depicting genes with greater than 1.5-fold change in gene expression (q<0.05). Genes upregulated/downregulated in lactate at acidic pH were directly compared to genes upregulated/downregulated in propionate. D) Venn diagrams depicting genes with greater than 1.5-fold change in gene expression (q<0.05). Genes upregulated/downregulated in the *phoT::Tn* strain compared to WT at pH 5.7 in lactate were compared to genes upregulated/downregulated in *phoR::Tn* at pH 5.7 in propionate.

Next, we examined genes associated with growth vs. growth arrest on each carbon source comparing *phoT::Tn* mutant at pH 5.7 in lactate to WT at pH 5.7 in lactate, as well as the *phoR::Tn* mutant at pH 5.7 in propionate vs. WT at pH 5.7 in propionate. This comparison allows us to investigate growth associated transcriptional regulation in each carbon source. There was a lack of overlap between upregulated genes (Fig. 7D), showing that suppressing growth arrest on lactate and propionate involve different processes. Many of the genes upregulated only in the *phoT::Tn* mutant are SenX3-RegX3 regulated, including *mgtA, udgA, pstA1, pstC2*, and *pe19* (Fig. 7B). Notably genes related to the electron transport chain are absent in this list. These data suggest that the *phoT::Tn* mutant is still experiencing dysregulation of the ETC caused by lactate.

We next examined genes that are differentially regulated by *phoT::Tn* compared to the WT at pH 5.7 as well as *phoT::Tn* compared to WT at pH 7.0. This comparison drove our understanding of the PhoT-regulated genes independent of pH. We observed a narrow overlap of gene regulation when making this comparison (Fig. 8D), suggesting limited pH-independent regulatory functions of *phoT*. The most notable overlaps of genes differentially expressed by *phoT::Tn* regardless of pH were a global upregulation of phosphate-related genes, an increase in SenX3-RegX3 regulated genes, an increase in ESX-5 substrate genes, and a decrease in PDIM genes (Fig. 8A-8C). This led to the conclusion that SenX3-RegX3 is activated in the *phoT::Tn* mutant, causing upregulation of phosphate transport machinery to compensate. When comparing WT at pH 5.7 to 7.0 and the *phoT::Tn* mutant at pH 5.7 vs. 7.0 (Fig. S10) we also observed that the PhoPR regulon was induced in both the WT and *phoT::Tn* mutant at acidic pH. Both the *phoT::Tn* mutant and WT demonstrate an upregulation of phosphate related genes. The ESAT-6 like protein locus *esxI-esxJ* is also upregulated in the *phoT::Tn* mutant regardless of pH (Fig. S10). Overall, the transcriptional signatures point towards the two-component system SenX3-RegX3, ESX-5, and the ETC being modulated during acid growth arrest on lactate.

**Figure 8.**
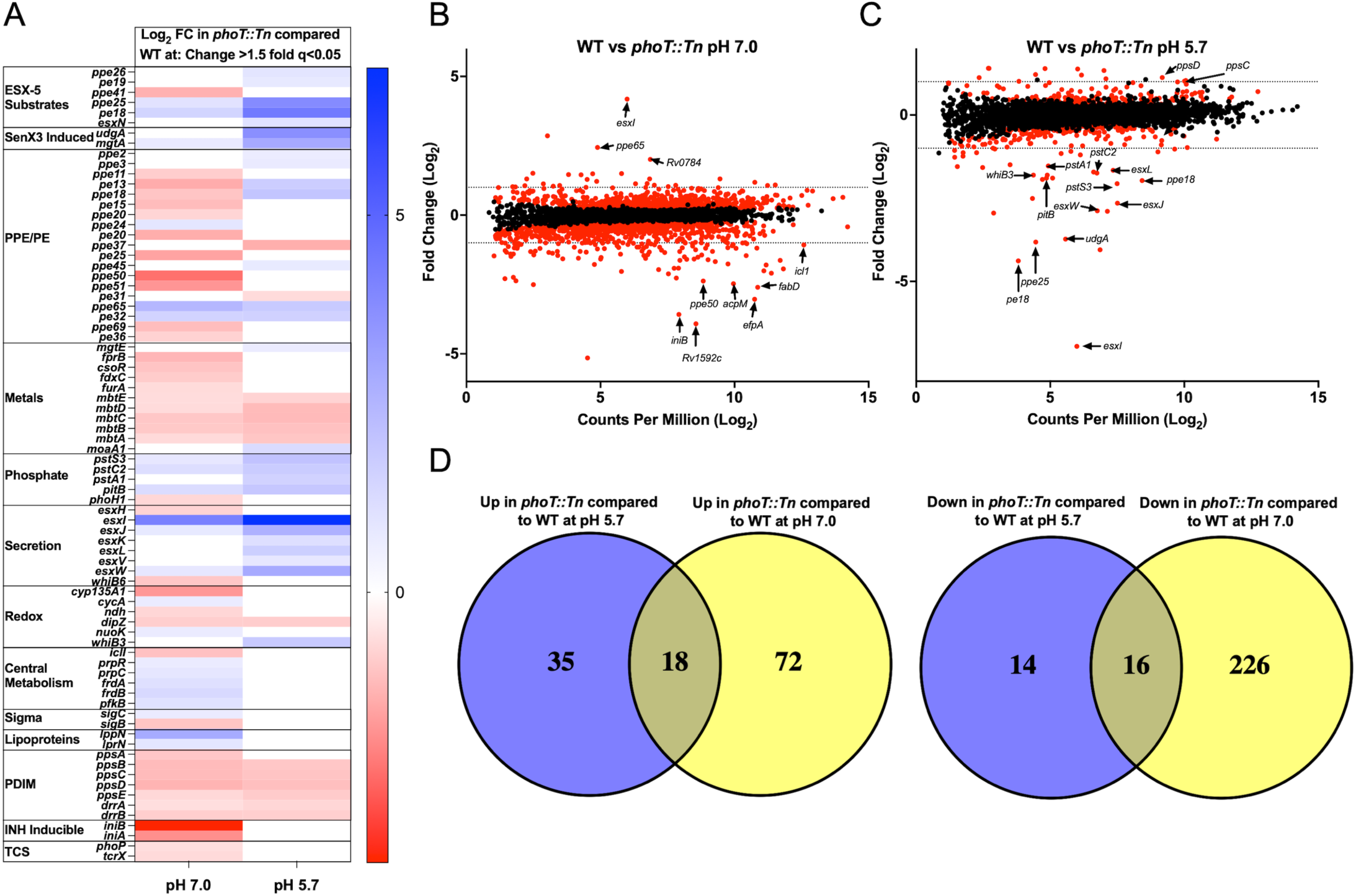
Differential gene regulation in the *phoT::Tn* mutant compared to the WT. A) Heat map of genes differentially regulated (fold-change>1.5, q<0.05) in the *phoT::Tn* mutant compared to WT at pH 7.0 (left) and pH 5.7 (right). Upregulation in the *phoT::Tn* mutant is indicated by blue and downregulation in the mutant is indicated by red. B) Magnitude/amplitude plots of average Log_2_ counts per million (CPM) and Log_2_ fold-change of the *phoT::Tn* mutant grown at pH 7.0 compared to the WT grown at pH 7.0. C) Magnitude/amplitude plots of average Log_2_ counts per million (CPM) and Log_2_ fold-change of the *phoT::Tn* mutant grown at pH 5.7 compared to the WT grown at pH 5.7. D) Venn diagram depicting the overlap of genes upregulated in the *phoT::Tn* mutant compared to the WT at both pH 5.7 and 7.0. Central region indicates genes upregulated/downregulated in the *phoT::Tn* mutant at both neutral and acidic pH compared to the WT. The blue region indicates genes only upregulated in the *phoT::Tn* mutant at acidic pH. The yellow region indicates genes only upregulated in the *phoT::Tn* mutant at neutral pH.

#### Growth rescue on lactate is dependent on the two-component regulatory system SenX3-RegX3

SenX3-RegX3 is a two-component regulatory system in Mtb where SenX3 is the membrane bound histidine kinase and RegX3 is the response regulator (41–43). A high phosphate concentration inhibits SenX3 autophosphorylation; under low phosphate conditions, inhibition is lifted and SenX3 autophosphorylates and phosphorylates RegX3 which subsequently regulates gene expression related to phosphate starvation and other pathways (41, 43, 44). The transcriptional profiling data strongly suggest that SenX3-RegX3 plays a role in growth arrest on lactate. Using a Δ*phoT* mutant and strains we previously generated (16, 45–47) we examined if the phosphate, lactate, and pH-dependent phenotypes were dependent on PhoT, PstA1, PE19, and RegX3. We generated a Δ*phoT* mutant (Fig S11) and complemented strain. In acidified medium with lactate as a sole carbon source, WT and complemented strains grew in the absence of phosphate but did not grow in the presence of phosphate (Fig. 9A). In contrast, in the presence of phosphate, the *ΔphoT* mutant grew to an OD_600_ of ∼0.4 (Fig. 9A). When phosphate is removed, the *ΔphoT* mutant can no longer grow, possibly due to the lack of stored phosphate that could accumulate in the WT when the bacteria were initially grown on rich medium. We hypothesized that, since PhoT interacts with PstSCA transport system to drive translocation of phosphate, disruption of another component of this phosphate transport system would phenocopy the Δ*phoT* mutant in this assay. We performed the same assay with a Δ*pstA1* mutant(16). PstA1 is the transmembrane domain of the phosphate transport system. The Δ*pstA1* mutant did indeed phenocopy the Δ*phoT* mutant where Δ*pstA1* grew to an OD_600_ of ∼0.3 in the presence of phosphate and did not grow in the absence of phosphate (Fig. 9B). We conclude that the phosphate starvation and enhanced growth on lactate phenotype is dependent on the PhoT-PstSCA complex, a finding consistent with similar studies on glycerol in modified, carbon replete 7H9 medium (17).

**Figure 9.**
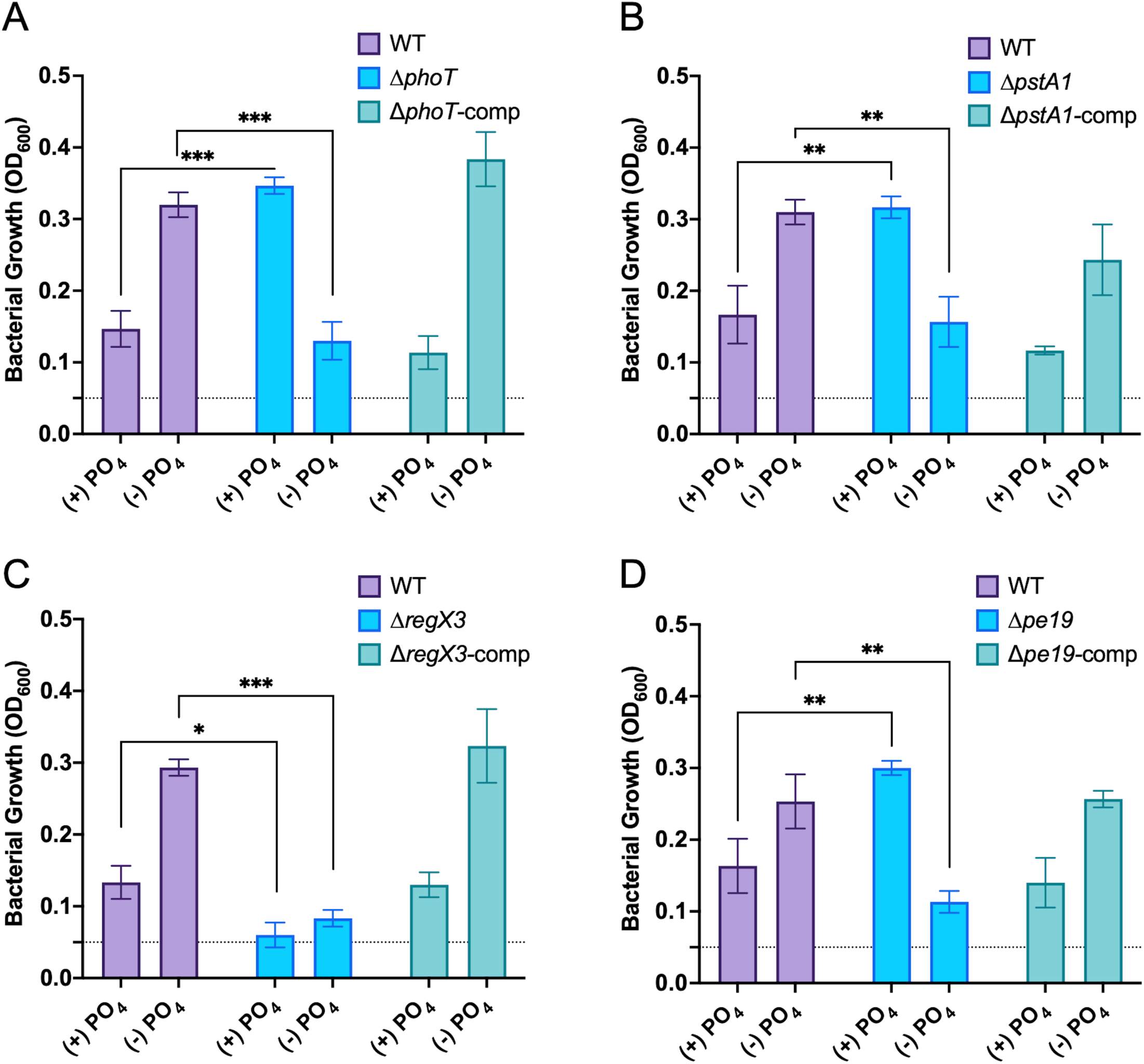
Growth on lactate is dependent on *phoT, pstA1, regX3, and pe19*. A) WT (purple), Δ*phoT* (light blue), and Δ*phoT* complemented strain (teal) grown in MMAT pH 5.7 either lacking phosphate completely or with the high 25 mM concentrations of phosphate in MMAT, 4 mM lactate was used. Statistical significance was determined using an unpaired t-test (***, p<0.001). B) WT (purple), Δ*pstA1* (light blue), and Δ*pstA1* complemented strain (teal) Similar medium described above. Statistical significance was determined using an unpaired t-test (**, p<0.005). C) WT (purple), Δ*regX3* (light blue), and Δ*regX3* complemented strain (teal) grown in similar medium as described above.. Statistical significance was determined using an unpaired t-test (*, p<0.05 ***, p<0.001). D) WT (purple), Δ*pe19* (light blue), and Δ*pe19* complemented strain (teal) in similar medium as described above. Statistical significance was determined using an unpaired t-test (**, p<0.005). All four experiments were performed at least twice with similar results. The error bars represent standard deviations of three technical replicates.

We hypothesized that the phosphate starvation sensing two-component system SenX3-RegX3 plays a role in the phosphate starvation dependent growth on lactate. To evaluate the role of SenX3-RegX3 on lactate-dependent acid growth arrest, we examined the WT, Δ*regX3*, and complemented strains in a similar experiment as described with the Δ*phoT* and Δ*pstA1* mutant strains. The Δ*regX3* strain could not grow on lactate in the presence or absence of phosphate (Fig. 9C) indicating that enhanced acid growth on lactate in phosphate-limited conditions is dependent on SenX3-RegX3. Lastly, PPE and PE proteins have recently been implicated in nutrient uptake (12, 48–51), with evidence for PE19 playing a specific role in phosphate-dependent growth arrest on glycerol (17). We therefore hypothesized that PE19, a PE protein in the SenX3-RegX3 regulon (47), may be associated with growth on lactate. We examined the growth of WT, Δ*pe19* and complemented strain and found that the Δ*pe19* mutant has a similar phenotype to the Δ*phoT* and Δ*pstA1* mutants, growing to an OD_600_ of approximately ∼0.3 in the presence of phosphate but not in the absence of phosphate (Fig. 9D). These data indicate that PE19 may also regulate phosphate sensing or uptake pathways to promote growth arrest at acidic pH on lactate.

#### Phosphate limitation promotes growth on glycerol, glucose, and lactate

A recently published study demonstrated that a Δ*phoT* strain could grow on glycerol at a pH as low as 5.0 (17). These data are in contrast to ours that show a *phoT::Tn* mutant cannot grow on glycerol at pH 5.7. Several differences exist between the studies, including the Mtb strains (Erdman vs. H37Rv) and the media used (MMAT vs. carbon depleted 7H9). Given the major differences in the media tested, where our MMAT medium is less complex than 7H9, we conducted studies using the media as described in Healy *et al* (*17*), with our strains and growth conditions. By changing the medium, we replicated the finding of Healy *et al*, observing robust growth of the Δ*phoT* and *phoT::Tn* strains on glycerol but no growth on glucose (Fig. S12A, S12B). These data indicate that a component of the medium present in carbon-depleted 7H9 that is absent from our MMAT medium permits growth of *phoT* mutant strains on glycerol. Lastly, we were curious if there were differences in growth profiles of WT on different non-permissive carbon sources when phosphate is depleted from MMAT medium, as compared to a mutation in *phoT*. We observed enhanced growth of WT on glucose, glycerol, and lactate (Fig. S12C) at low phosphate. The differences between growth of the *phoT::Tn* mutant (Fig. 2A) and phosphate depleted WT (Fig. S12C) on non-permissive carbon sources shows that the *phoT::Tn* mutant does not phenotypically replicate a low phosphate environment, and suggests more complex interactions of pH and phosphate are at play, controlling growth on specific carbon sources.

## Discussion

Mtb establishes a state of non-replicating persistence on lactate at acidic pH, and this growth arrest phenotype can be overcome by genetic mutation. This finding further supports our “acid fasting” hypothesis that growth arrest at acidic pH is due to metabolic restrictions (9). Growth arrest on glycerol is overcome by increasing uptake of glycerol through dominant mutations in *ppe51* (12), growth arrest on propionate is overcome by loss-of-function mutations in *phoPR* resulting in rerouting carbon from lipid synthesis to central metabolism (13), and as described here, growth arrest on lactate is overcome by limiting phosphate. The three distinct mechanisms of growth arrest support that Mtb has evolved to constrain its growth in acidic environments, and that availability of specific carbon sources and other nutrients such as phosphate can regulate persistence versus growth.

Mtb encounters acidic pH when it is phagocytosed by a macrophage and also possibly in some granulomatous tissues, where pH of the granuloma has been measured in the pH 5.0 to pH 7.2 range (52). In the guinea pig Mtb infection model, lactate accumulates in granulomatous tissues as infection proceeds (53). Stanley *et al*, 2024, demonstrated that ongoing accumulation of mutations and evolution of the lactate dehydrogenase gene *lldD2* in tuberculosis is associated with a selective advantage in adjusting to host associated stresses (54). It has also been shown that Mtb infection in mouse lungs induces the Warburg Effect wherein inflamed tissue around the infected area go through aerobic glycolysis to produce lactate (55–57). Additionally, lactate concentrations in lung tissue within a *Staphylococcus aureus* infected rabbit can reach as high as 15 mM (58). L-lactate is the primary isomer in the host, which is the isomer used for this study. However, a recent publication found that Rv1257c may be acting as a D-lactate dehydrogenase, implying it may be present for Mtb to utilize in the host. The same study also demonstrated that a *ppe3* transposon mutant had a growth defect *in vitro* on both L- and D- lactate (59). It has also been widely reported that lactate acts as an immunomodulatory signaling molecule that can act in both autocrine and paracrine fashions (60–62). Ultimately, the experimental difficulty of measuring the metabolic contents of a phagosome during active infection means there is a knowledge gap in the availability of lactate and phosphate to Mtb replicating in a phagosome. However, due to the abundance of lactate in the surrounding tissue during infection, and the fact that the phagosome is an environment where phosphate may be limited, we predict that this combination of signals may be present during active infection, especially relevant to extracellular Mtb in necrotic tissue. Integration of lactate concentration, phosphate availability, and pH may provide information to the bacterium about nutrient availability and the immune environment to regulate growth.

The impact of lactate on bacterial physiology has been well documented due to its relevance to industrial processes (63–65). We predicted that growth arrest on lactate may be due to similar stresses imparted on lactic acid bacteria by the excessive production and accumulation of lactate. These stressors include but are not limited to: acidification of the cytoplasm (21), decreases in membrane potential by permeabilizing the membrane (22), and cytoplasmic acidification that decreases proton motive force (PMF). Mtb however has been shown to tightly regulate its cytosolic pH (5, 8, 66), and any deviation from the typical ∼7.2-7.5 can be indicative of stress. The ETC is intimately linked with PMF, where it maintains the proton gradient for ATP synthesis as well as producing reduced cofactors. In this study we demonstrate that acidic pH, lactate and phosphate act together to reduce PMF, which may drive growth arrest at acidic pH.

We propose a two-part, stress and nutrient uptake model to explain Mtb growth arrest on lactate at acidic pH. In the first part, the cytoplasmic pH stress model, we hypothesize that the combined impacts of lactate and phosphate place physiological constraints on the bacterium as it adapts to maintain pH-homeostasis and PMF needed for oxidative phosphorylation or maintain protein folding. In this model, when either lactate or phosphate enter the cell, the proton associated with each molecule disassociates and is released into the cytoplasm, causing cytoplasmic acidification (Fig. 4-5). When available as a sole carbon source, lactate drives Mtb to pump out protons to maintain pH-homeostasis, resulting in increased membrane potential. Inactivating mutations of *phoT* through transposon insertion decrease the amount of phosphate that is entering the cell, alleviates some of the cytoplasmic acidification stress and allows the bacteria to maintain a higher membrane potential. The combination of a more neutral cytoplasmic pH and higher membrane potential leads to a PMF that is sufficient for oxidative phosphorylation and growth. We speculate that when phosphate and lactate surpass specific thresholds, Mtb cytoplasmic pH drops below a threshold of pH 6.8, where the PMF is insufficient to support oxidative phosphorylation. This model is supported by our RNA-sequencing data in which we observed upregulation of ETC associated genes under lactate at acidic pH but not propionate. It is also possible that at lower ends of the observed cytoplasmic acidification (e.g. pH 6.6), that key metabolic enzymes may unfold, resulting in blocks of metabolic pathways (20, 67–72). An interesting observation from the RNA-sequencing data was that the arginine biosynthesis pathway was upregulated. In *E. coli*, arginine can be used as a proton sink in response to cytoplasmic acid stress(73). Additionally, to mitigate acid stress, *E. coli* can decarboxylate arginine, consuming a proton and producing agmatine that is exported out of the cell (73). Altogether, these data suggest that acidic pH accompanied by lactate and phosphate uptake cause cytoplasmic acidification and constrain growth. This cytoplasmic pH acidification can be mitigated by altering relative abundances of lactate, phosphate or extracellular pH.

In the second part, we hypothesize that when phosphate uptake is restricted, through *phoT* mutation or phosphate limitation, SenX3-RegX3 is induced, upregulating ESX-5 genes which aid in exporting of the PPE/PE proteins that may remodel the cell envelope or promote uptake of other nutrients (Fig. 10). In this model, limiting intracellular phosphate concentrations either through a mutation in a phosphate transporter (*pstSCA-phoT* system) or limiting extracellular availability, triggers the SenX3-RegX3 two-component system. (41, 43, 44). Tischler *et al* 2013 demonstrated that the deletion of the *pstA1* transmembrane domain of the phosphate transport system PstSCAB constitutively activates SenX3-RegX3 signaling (16). Since PhoT is an orphan ATPase that putatively interacts with PstSCA, specifically PstS3, PstC2, and PstA1, we propose a similar model, where mutating the ATPase subunit of this phosphate transport system also constitutively activates SenX3-RegX3 signaling. The SenX3-RegX3 regulon has been well characterized, as it regulates genes associated with phosphate (16, 41, 74), and activates the redox sensor WhiB3 (75) which is upregulated in our transcriptional profiling data (Fig. 6A, S10A). Mutating *pstA1* constitutively activates SenX3-RegX3 leading to constitutive expression of *udgA*, *mgtA*, and many other genes (16, 45, 47) which we also observe in the *phoT* mutant at both pH 5.7 and pH 7.0 (Fig. 8A). Based on our data, we conclude that a Δ*phoT* phenocopies the Δ*pstA1* strain. Due to the lack of growth of the Δ*regX3* mutant on lactate either with or without phosphate, we conclude that growth arrest on lactate can be overcome by the activation of SenX3-RegX3. Given that SenX3-RegX3 regulates a large number of genes, we speculate that some gene(s) downstream of SenX3-RegX3 signaling permits growth on lactate, through alleviating the pH associated stress with lactate or possibly promoting cytoplasmic pH and PMF homeostasis. Alternatively, a PE/PPE protein in the SenX3-RegX3 regulon may enable uptake of a nutrient that alleviates pH associated stress, such as magnesium or other metals in the medium. Lastly, given that the Δ*pe19* mutant phenocopies the Δ*phoT* and Δ*pstA1* mutants (Fig. 9D), this suggests it may be associated with transport of phosphate or some other nutrient necessary for growth. Ultimately these results show that SenX3-RegX3 can be triggered by either knocking out *phoT* or *pstA1*, and that potentially a protein regulated by SenX3-RegX3 allows growth on lactate at acidic pH.

**Figure 10:**
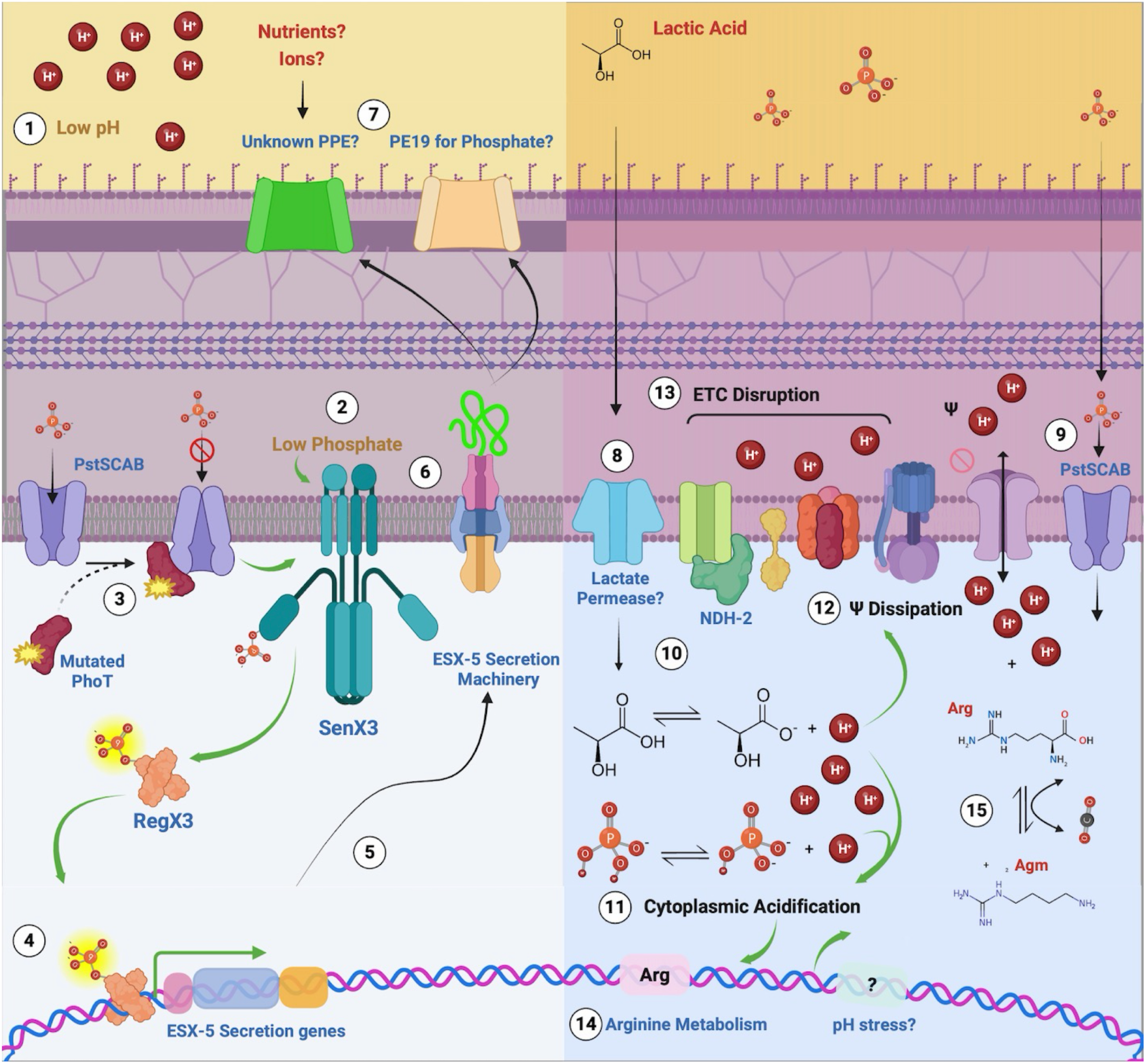
Proposed model for the impact of lactate and phosphate on Mtb growth at acidic pH. At acidic pH (∼5.7) (1), Mtb tightly regulates cytoplasmic pH to around 7.2-7.4. Under low phosphate conditions (2) or failure to uptake phosphate (3) (i.e., resulting from a mutation in any component of the phosphate transporter PstSCA, or its associated orphan ATPase, PhoT), the two-component regulatory system SenX3-RegX3 signaling is activated, resulting in the phosphorylation of the response regulator RegX3 (4), and the expression of genes in its regulon, including the components of the ESX-5 secretion system. Some unknown genes in the RegX3 regulon might also be involved in alleviating the pH stress induced by acidic growth conditions or an agent acidifying the cytoplasm. The ESX-5 system is assembled and inserted into the cytoplasmic membrane (5), where it exports several proteins to the mycomembrane (6), some of which can be PPE and PE proteins (e.g., PE19), which may play a role in nutrient acquisition and transport, regulation, or ion balance through the acquisition of certain nutrients or metal ions (7). A putative permease can then transport lactate from the pseudoperiplasmic space into the cytoplasm (8), which, when coupled with the transport of phosphate (9), dissociate into their ionized form, releasing protons (10), overwhelming the pH homeostasis and acidifying the cytoplasm of the bacteria (11). Additionally, maintenance of cytoplasmic pH-homeostasis on lactate, results in proton export modestly increasing membrane potential (!) (12), which is also increased by loss of phosphate import. Since cytoplasmic pH establishes the proton gradient required for the electron-transport chain (ETC) to function properly, cytoplasmic acidification lowers PMF and disrupts the ETC (13). As a countermeasure to cytoplasmic acidification, Mtb upregulates genes responsible for the synthesis of arginine (14), an amino acid that can act as a sink for the excess protons among other genes (15). Green arrows indicate positive influence (i.e., increase signaling or expression).

We previously performed RNA-seq analysis under similar conditions presented here, but with WT CDC1551 Mtb at acidic pH and neutral pH when exposed to either pyruvate or glycerol (8), allowing us to further define acidic-pH dependent adaptations that are lactate specific. The limitation of this comparison is that the lactate data was generated in the Erdman strain while glycerol and pyruvate were generated in the CDC1551 strain. When comparing WT Mtb grown in either pyruvate, glycerol, or lactate at acidic pH compared to neutral pH, 292 of the 529 genes differentially upregulated in lactate are specific to lactate at acidic pH (Fig. S13). This result supports the conclusion that growth arrest on lactate is mechanistically different than growth arrest on glycerol or pyruvate. When looking at the genes specifically upregulated under lactate at acidic pH, reinforcing patterns emerge. Cytochrome biogenesis genes *ccdA*, *ccsA*, *ctaE*, and *qcrCAB*, as well as the phosphate transport related genes *pstS3*, *pstC2*, and *pstA1* are only upregulated in lactate at acidic pH relative to neutral pH and not under glycerol or pyruvate at acidic pH. This is another potential indicator that lactate is causing PMF stress requiring upregulation of cytochrome biogenesis genes, and that phosphate transport and signaling is strongly related to lactate metabolism. This fits our model and hypothesis that lactate impacts PMF. In *Bacillus subtilis* cytochrome bd expression is linked to lactate dehydrogenase expression under microaerobic growth (76). Noticeably, the Region of Difference 1 (RD1), which is important for infection (77), is downregulated only under lactate at acidic pH but not pyruvate and glycerol. These data suggest that different virulence factors potentially are differentially expressed under different carbon sources. Altogether these data reinforce our hypotheses that Mtb uses a combination of acidic pH and carbon source availability to modulate genes important for the ETC, phosphate transport, metal regulation, and virulence.

Concurrent with our study, Healy *et al,* demonstrated that Δ*phoT* and Δ*pstA1* mutants are capable of growing on glycerol at acidic pH (17). Although we did not observe this phenotype in our studies with our *phoT::Tn* mutant (Fig. 2A) in MMAT medium, we were able to replicate their data by using modified 7H9 medium (17). Substantial differences exist between MMAT and 7H9, and it is important to determine which differences in the media drive specificity for lactate in MMAT and the *phoT::Tn* mutant. Additionally, we could grow WT Erdman at pH 5.7 in glycerol simply by depleting phosphate from our MMAT medium, suggesting that the inability of the *phoT::Tn* mutant to grow on glycerol is potentially due to a hypomorphic phenotype, where residual phosphate taken up by PstSCA-PhoT restricts growth on glycerol but not lactate. Understanding the nuances of these complex interactions should provide deeper insights into mechanisms that fine tune growth on specific carbon sources at acidic pH.

## Materials and Methods

### Bacterial strains and media

Experiments were performed using Mtb Erdman and grown at 37°C in standing, filter-vented T25 or T75 tissue culture flasks unless otherwise specified. Cultures were maintained in 7H9 Media supplemented with 10% OADC, 0.5% glycerol, and 0.05% Tween-80. Experiments performed in minimal defined MMAT medium as described by Lee *et al*, 2013: 1 g l^−1^ KH_2_PO_4_, 2.5 g l^−1^ Na_2_PO_4_, 0.5 g L^−1^ (NH_4_)_2_SO_4_, 0.15 g L^−1^ asparagine, 10 mg l^−1^ MgSO_4_, 50 mg L^−1^ ferric ammonium citrate, 0.1 mg L^−1^ ZnSO_4_, 0.5 mg L^−1^ CaCl_2_ and 0.05% Tyloxapol. The media was buffered to pH 5.7 of 7.0 using 100 mM MES or MOPS respectively (78). In the media lacking phosphate, KH_2_PO_4_ and Na_2_PO_4_ were omitted, then supplemented back in the form of KH_2_PO_4_. To replicate the Healy et al medium, we made 7H9, 10% Albumin, Sodium chloride using fatty acid free albumin, buffered to pH 5.0 using MES. 0.2% glycerol or 10% dextrose was then supplemented back for the two different mediums. In all experiments, cultures were grown to mid-log phase, then washed then seeded in experimental growth medium (MMAT high phosphate, MMAT low phosphate, or modified 7H9) at an OD_600_ of 0.05.

The Mtb Erdman Δ*pstA1*, Δ*pstA1*-comp, Δ*regX3*, Δ*regX3*-comp, Δ*pe19* and Δ*pe19*-comp strains were previously described(16, 47). The Mtb Erdman Δ*phoT* mutant was made by allelic exchange with pJG1100-Δ*phoT*. Regions of the *Mtb* Erdman genome ∼850 bp 5’ and 3’ of *phoT* were amplified by PCR and cloned in pJG1100 between the PacI and AscI sites and Sanger sequenced. The pJG1100-Δ*phoT* plasmid was electroporated in *Mtb* Erdman and transformants were selected on 7H10 agar with kanamycin (15 μg/ml) and hygromycin (50 μg/ml). Transformants were then grown without antibiotics and plated on 7H10 with 2% sucrose to counterselect the pJG1100 plasmid as described(16). The Δ*phoT* mutant was confirmed by PCR and Southern blotting. Production of the virulence lipid phthiocerol dimycocerosate (PDIM) by the Δ*phoT* mutant was verified thin layer chromatography of ^14^C propionate-labeled cell wall lipids as described(16). Primer sequences are available upon request.

The complementation plasmids for the *phoT::Tn* strain were generated using a single-copy chromosomal insertion vector (pMV306) and an episomal overexpression vector (pVV16). The WT gene *phoT* was cloned into pMV306 using Gibson cloning, taking approximately ∼1000 bp upstream to include the native promoter. The complete pMV306::*phoT* construct was electroporated into *phoT::Tn* strains and selected for on 7H10 plates with zeocin (25 µg/mL). WT *phoT* was cloned into pVV16 using Gibson cloning. For the pVV16 construct, the *phoT* gene was cloned under the *hsp60* promoter. The complete pVV16::*phoT* construct was electroporated into *phoT::Tn* strains and selected on 7H10 agar with hygromycin (50 µg/mL).

### Construction of Transposon Library

A Mtb Erdman transposon library was generated using the φMycoMar T7 phage as previously described (79, 80). Wild type Mtb was grown to mid-log phase in 50 mL of 7H9 supplemented with 10% OADC and 0.05% Tween-80, after which bacteria were washed and incubated in 4.5 mL of MP buffer (50 mM Tris-HCl pH 7.5, 150 mM NaCl, 10 mM MgSO_4_, and 2 mM CaCl_2_) with 1x10^11^ PFU φMycoMar T7 at 37°C for 4 hours. The Mtb-phage mix was then split between 8 or more 15 cm 7H10 plates containing 25 µg/mL kanamycin and incubated for three to six weeks at 37°C. Libraries with totaling ∼100,000 CFUs were collected in 2-4 separate pools and stored at -80°C for subsequent use.

### Genetic selections

To select for spontaneous mutants, Mtb Erdman was grown to an OD_600_ of 0.6-1.0, spun down, and resuspended in MMAT media buffered to pH 5.7. Mtb cells were calculated for 10^9^ CFU and plated across 4 MMAT agar plates buffered to pH 5.7 supplemented with 4 mM lactate. Plates were incubated at 37°C for over 12 weeks without any isolated colonies appearing.

To select for transposon mutants, an Mtb Erdman transposon library consisting of >100,000 mutants were thawed and grown for 24 hours in 10 mL unbuffered 7H9 supplemented with 10% OADC and 0.05% Tween-80 to allow for initial growth. The OD_600_ was measured on a spectrophotometer, calculated for 10^9^ CFU, and plated across 4 MMAT agar plates buffered at pH 5.7 and amended with 4 mM lactate. Plates were incubated for 4-8 weeks at 37°C after which 20 isolated colonies were randomly chosen for follow-up growth studies. Isolated colonies were confirmed for growth at acidic pH in liquid culture (MMAT + 4 mM lactate buffered to pH 5.7). The inverse PCR technique (18) was used initially on two colonies to identify transposon insertion sites but was later replaced using whole genome sequencing (WGS) combined with the BLAST-like alignment tool (BLAST) (81) for more robust results. For WGS, genomic DNA was extracted and sequenced using the Illumina MiSeq and a 200 Mbp (1.33M reads) sequencing package. Following the run, reads were trimmed and the variant caller breseq (82) was used to align and compare sequencing data against the Erdman or CDC1551 reference genomes. BLAST was used to map the φMycoMar T7 transposon sequence onto assembled reads and find the transposon insertion breakpoints. The NCBI reference number for the mapping of the Erdman Tn insertion sites is NZ_KK339487.1.

### Lactate-and-phosphate dose response checkerboard assay

The checkerboard experiment was performed in black-sided, clear bottom 96-well plates. Medium was seeded with Mtb at an OD_600_ of 0.05 in wells with a 2.5-fold dilution series KH_2_PO_4_ (0 μM – 200 μM) and a 2.5-fold dilution series of lactate (0 μM – 40000 μM) examining all combinations. The plates were then incubated for 21 days. Before taking readings in the plate reader, the cells were resuspended to ensure homogeneity. ODs were normalized to percent-of-maximum using wild-type minimum and maximum growth values. Each condition was conducted in triplicate.

### Cytoplasmic pH assay

The pH of the cytoplasm was measured using the protocol described by Purdy *et al.*, using the pH-sensitive dye CMFDA (83). Mtb was grown in rich 7H9 to mid-log phase, cells were pelleted and resuspended in fresh 7H9 in 500 µL at an OD600 of 0.5. 4 μL of 1 μg mL^-1^ CMFDA dye was added to each sample and incubated for 2 hours at 37°C. Samples were then washed twice in 7H9 to remove excess CMFDA. After the last wash, the cells are resuspended in MMAT with phosphate omitted, then transferred to a 96-well plate where lactate and phosphate were added, fluorescence was then measured with excitations of 450 nm and 490 nm and emission at 520 nm for the course of 60 minutes. A standard curve was generated using Mtb in buffered medium treated with 10 μM nigericin.

### Measuring membrane potential

Mtb was grown in rich 7H9 to mid-log phase, cells were pelleted and resuspended in fresh 7H9 in 500 µL at an OD600 of 0.5. The fluorescent dye DiOC_2_ was added to a final concentration of 30 µM. Cells were labeled for 30 minutes at room temperature. The cells were then washed twice in 7H9 to remove excess DiOC_2_. After the final wash, cells were resuspended in MMAT without phosphate then transferred to a 96-well plate. Carbonyl cyanide m-chlorophenyl hydrazone (CCCP) was used as a depolarizing agent at a final concentration of 50 µM., Emission spectra was read at 610 and 515 nm after exciting at 485 nm over the course of 60 minutes.

### ETC inhibitor dose response assays

Mtb was grown up to mid-log phase, cells were washed and then seeded in a black sided, clear bottom 96 well plate at an OD of 0.05. Plates were incubated at 37°C in a plastic bag containing a wet paper towel to maintain humidity and prevent evaporation. Plates were read after 10 days of incubation. For each strain a rifampicin and DMSO treatment were included as positive and negative controls that were later used for normalization.

### Transcriptional profiling and data analysis

Initial Mtb cultures (wild type, phoT::*Tn* mutant 2.4, and *phoR::Tn*) were grown in 7H9 buffered media at 37°C and 5% CO2 in T-25 vented flasks. Cultures were expanded out to 75 mL 7H9 buffered media. Once mid-log growth phase was reached a second time, cultures were spun down and pre-adapted for three days in MMAT (pH 7.0 + glycerol) to account for metabolite carryover. Experimental cultures were inoculated at a starting OD_600_ of 0.6 in 8 mL of MMAT buffered media amended with 4 mM lactate or 2 mM propionate. Treatment conditions examined include i) wild type Mtb at pH 5.7 with lactate, ii) wild type Mtb at pH 5.7 with propionate, iii) wild type Mtb at pH 7.0 with lactate, iv) wild type pH 7.0 with propionate, v) *phoT::Tn* at pH 5.7 with lactate, and vi) *phoT::Tn* at pH 7.0 with lactate, vii) *phoR::Tn* at pH 5.7 with propionate, viii) *phoR::Tn* at pH 7.0 with propionate. Each treatment condition was performed with two biological replicates and incubated for three days after which total bacterial RNA was extracted as previously described using the TRIzol method from the Qiagen RNeasy kit (ID: 74104) (8, 84). RNA quantity and quality was assessed by Qubit Fluorometer using the Qubit RNA BR kit (ID: Q10210) and Agilent 2100 Bioanalyzer, respectively. DNA was depleted using the Invitrogen TURBO DNA-free kit (ID: AM1907). Finally, the RNA library was sequenced using a NovaSeq 6000. Raw sequencing data was analyzed using SPARTA (ver. 1.0) (85). Genes were filtered based on > 1.5-fold differential gene expression and q < 0.05. Transcriptional profiles are deposited in the GEO Database (accession number GSE301034).

## Acknowledgements

We are grateful to Veronica Albrecht for technical assistance. This work was funded by the NIH-NIAID (R01 AI116605, R01 AI150855 and R01 AI173285) and AgBioResearch.

## Author contributions

Shelby Dechow – Conceptualization, investigation, and editing. Adam Kibiloski – Conceptualization, investigation, visualization and writing. Heather Murdoch – investigation and editing, Bassel Abdalla – investigation visualization and editing, Anna D. Tischler – investigation, resources and editing; Robert Abramovitch – Conceptualization, investigation, resources, supervision, writing, and editing.

## Disclosures

R.B.A. is the founder and owner of Tarn Biosciences, Inc., a company that is working to develop new antimycobacterial drugs.

---

**Figure S1.**
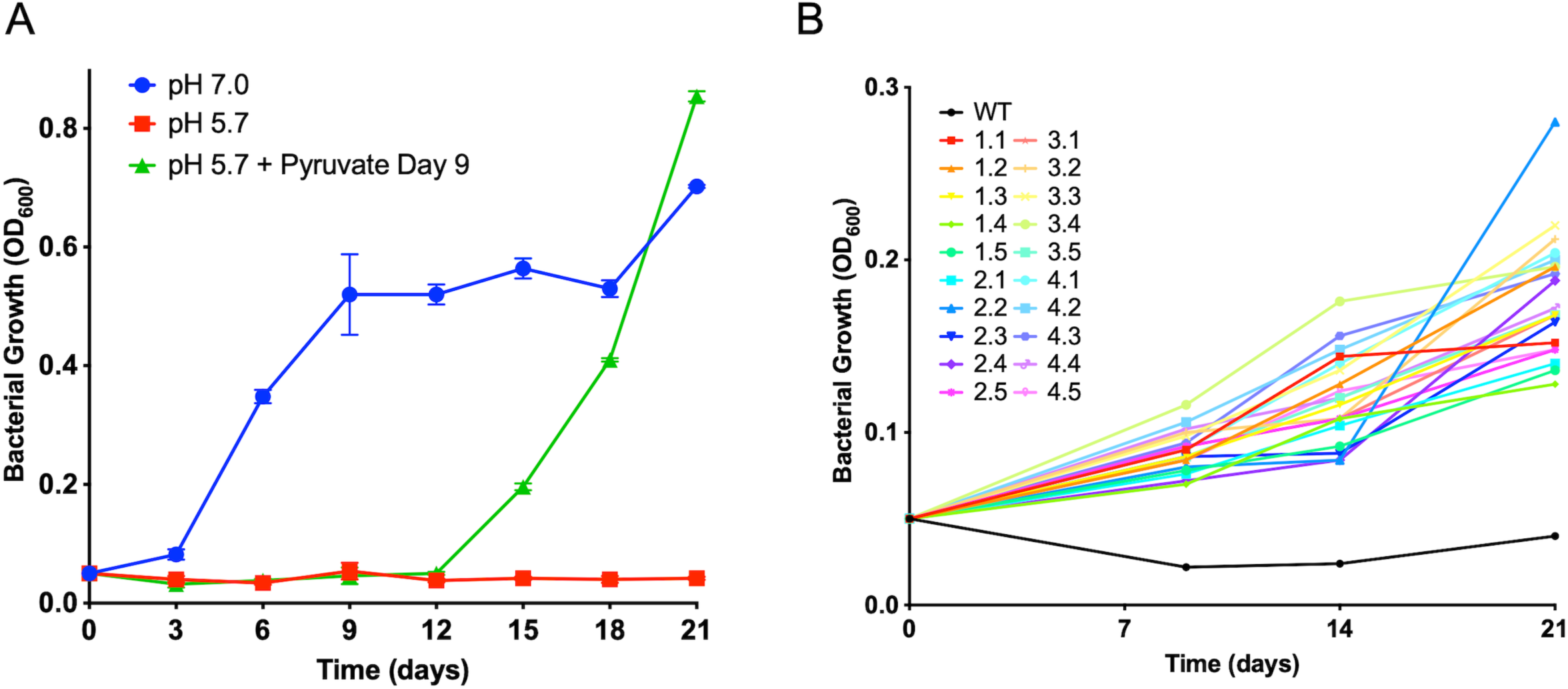
Growth arrest on lactate can be overcome through pyruvate supplementation or genetic mutation. A) WT CDC1551 was grown in minimal medium buffered to pH 5.7 (red) and 7.0 (blue) supplemented with 4 mM lactate. On day 9, 10 mM pyruvate was added to Mtb grown on lactate at pH 5.7 (green). This experiment was performed once in biological duplicates. B) Growth curves of the transposon mutants grown on 4 mM lactate at acidic pH 5.7 over 21 days, OD_600_ were measured on day 9, day 13 and day 21. This experiment was performed once in biological duplicate

**Figure S2.**
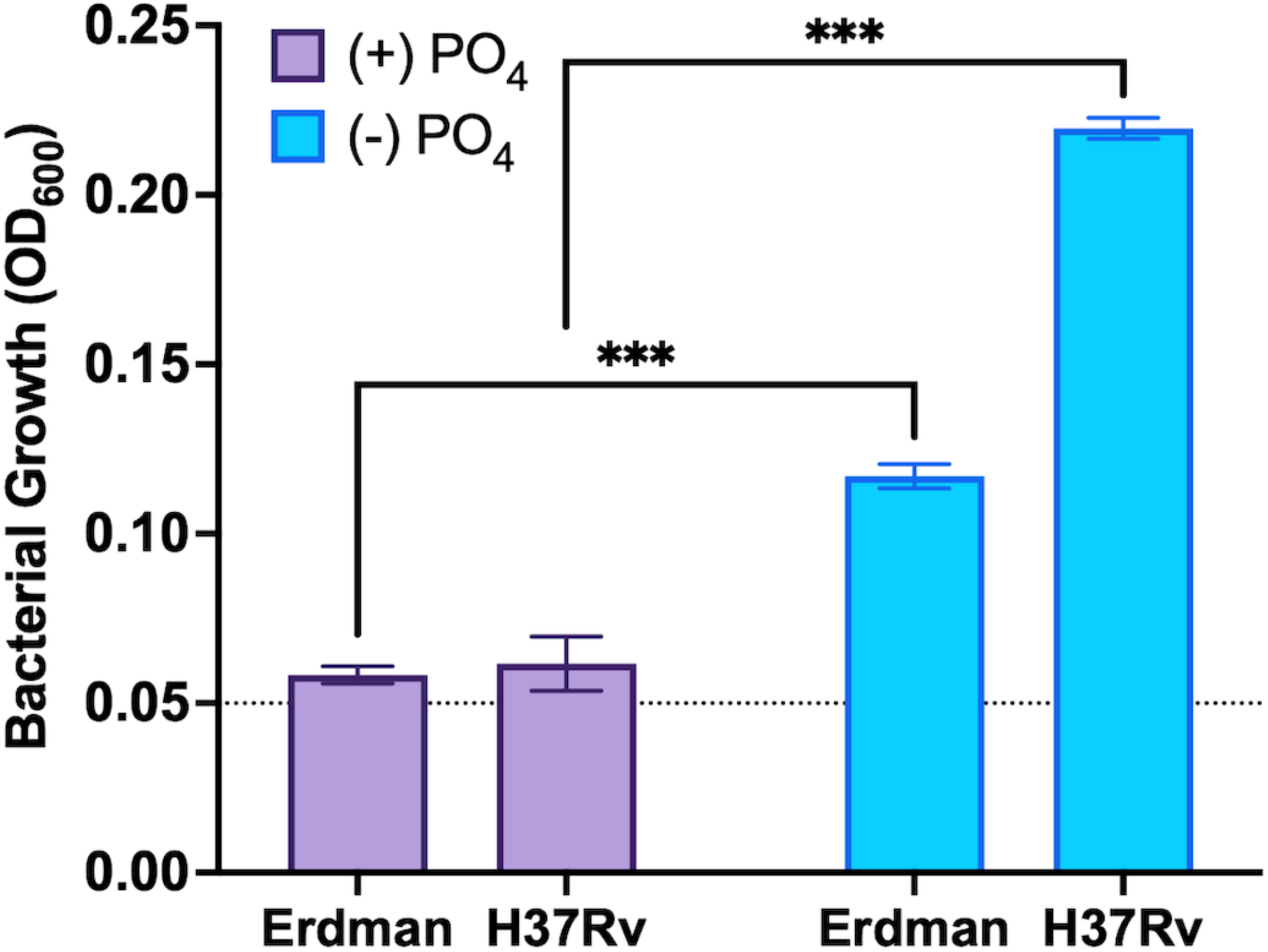
Growth arrest in lactate can be overcome in both WT H37Rv and WT Erdman by depleting phosphate. Growth of WT Erdman strain and WT H37Rv in medium supplemented with 4 mM lactate at pH 5.7 in medium either containing phosphate (purple) or not containing phosphate (blue) after 21 days. Dashed line indicates growth arrest phenotype. Statistical analysis was performed using an unpaired t-test (***, p<0.001). Experiment was performed twice. Error bars represent the standard deviation of three technical replicates.

**Figure S3.**
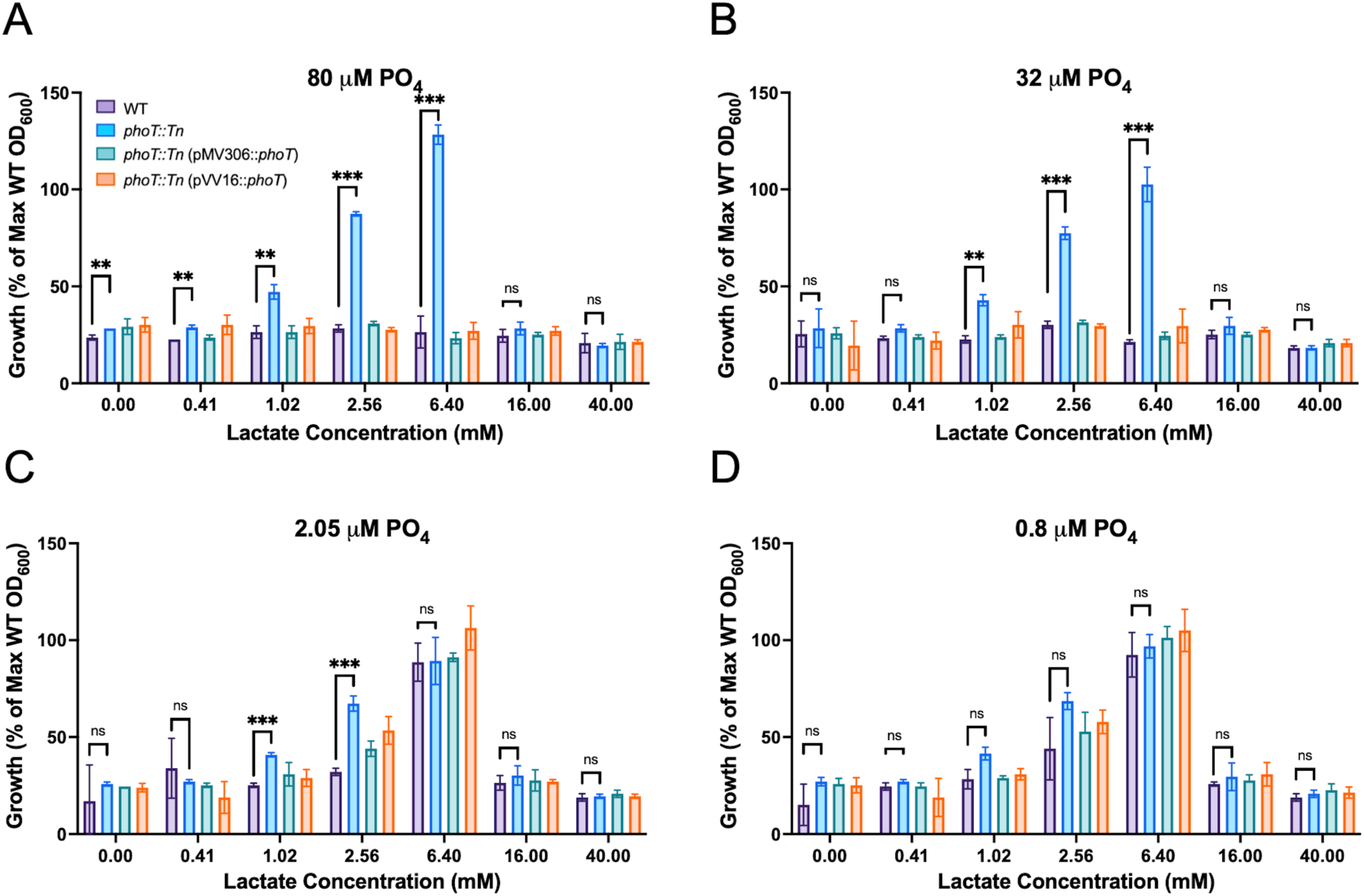
Mtb growth arrest on lactate is dependent on phosphate concentration. Mtb grown at varying lactate concentrations in media amended with (A) 80 µM, (B) 32 µM, (C) 2.05 µM, and (D) 0.8 µM phosphate. Statistical analyses were performed using an unpaired t-test (*, p<0.05 **, p<0.005 ***, p<0.001). The experiment was repeated twice with similar results. The error bars represent the standard deviation of three technical replicates. This experiment was performed at least twice. Error bars represent the standard deviation of three technical triplicates.

**Figure S4.**
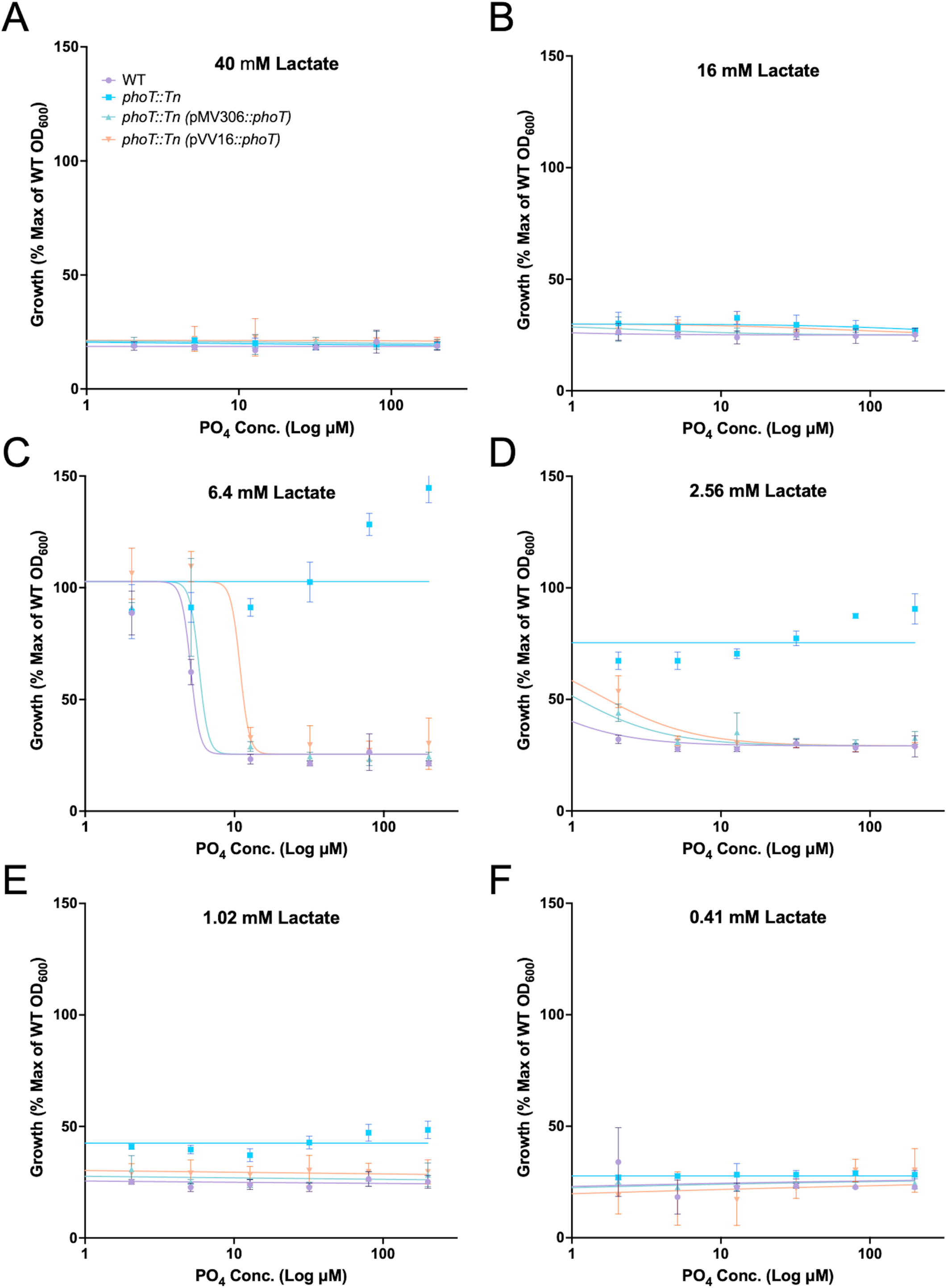
Growth arrest on lactate is dependent on phosphate concentration. Mtb grown at varying lactate concentrations in media amended with (A) 40 mM, (B) 16 mM, (C) 6.4 mM, (D) 2.56 mM, (E) 1.024 mM, (F) 0.41 mM lactate. Statistical analyses were performed using an unpaired t-test (***, p<0.001). This experiment was performed at least twice. Error bars represent the standard deviation of three technical triplicates.

**Figure S5.**
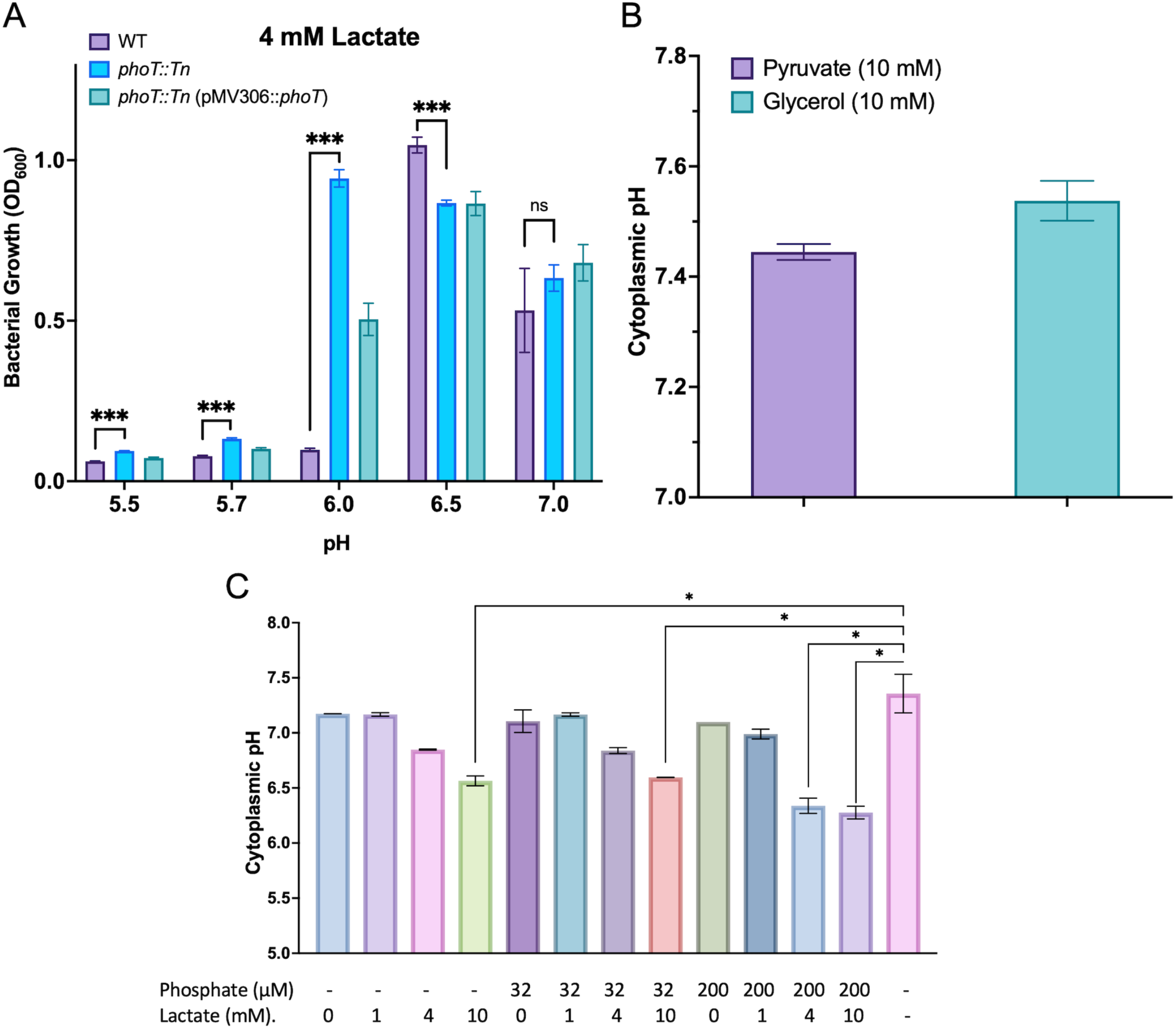
Growth arrest on lactate is pH dependent. A) Growth of WT (purple), *phoT::Tn* (light blue), and the complement (teal) was evaluated after 21 days. Statistical significance was evaluated using an unpaired t-test (***, p<0.001). This experiment was performed once. Error bars represent the standard deviation of three technical replicates. B) Cytoplasmic pH measurements of Mtb exposed to either pyruvate (purple) or glycerol (teal). This experiment was performed at least twice with similar results. Error bars represent the standard deviation of three technical replicates. C) Endpoint data from the cytoplasmic pH time course experiment where. All cytoplasmic pHs’ were compared to the WT DMSO control exposed to no lactate or phosphate. Statistical significance was evaluated using an unpaired t-test (*, p<0.05). This experiment was performed twice with similar results. Error bars represent the standard deviation of three technical replicates.

**Figure S6.**
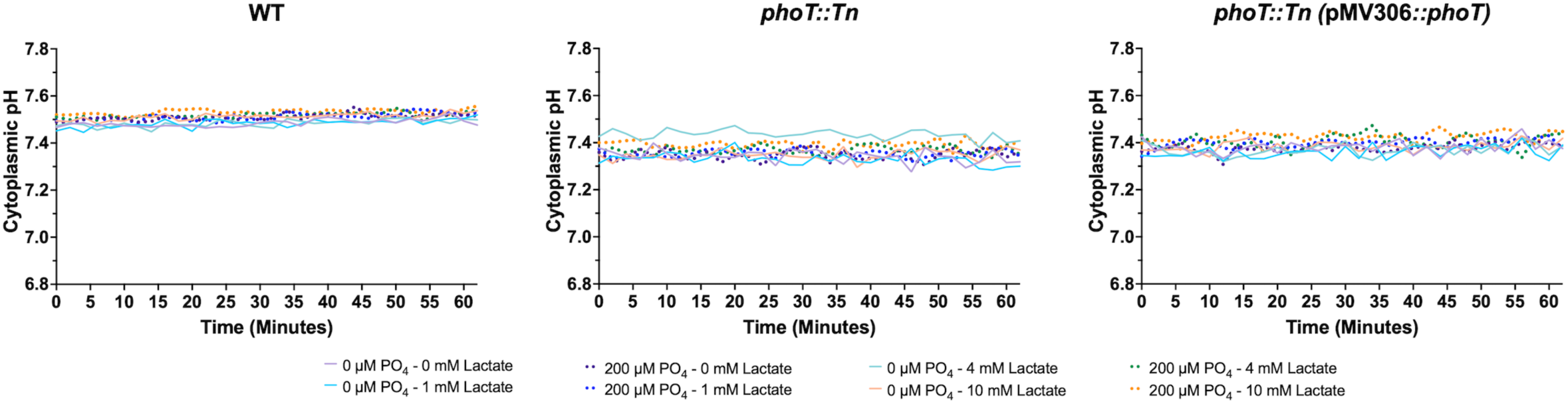
Cytoplasmic pH in not impacted in strains exposed to lactate and phosphate at neutral pH. WT*, phoT::Tn,* and complemented strains exposed to a range of concentrations of lactate and phosphate at neutral pH. Experiment were performed at least twice with similar results. Error bars represent the standard deviation of three technical triplicates.

**Figure S7.**
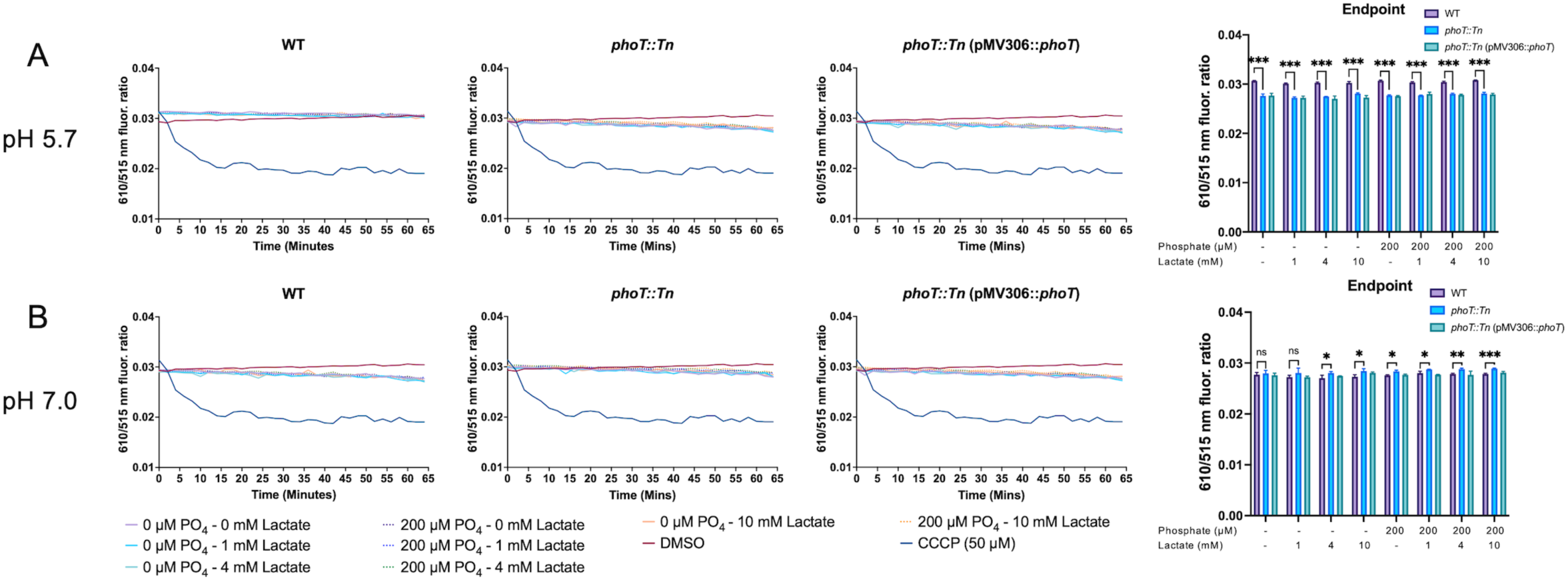
Lactate and phosphate at low concentrations have modest impacts on membrane potential. A) WT, *phoT::Tn*, and complemented strains membrane potential monitored over time at pH 5.7 with endpoint data plotted. Statistical significance determined by unpaired t-test (***, p<0.001). B) WT, *phoT::Tn*, and complemented strains membrane potential monitored over time at pH 7.0 with endpoint data plotted. Statistical significance determined by unpaired t-test (*, p<0.05, **, p<0.005, ***, p<0.001). Both experiments were performed at least twice with similar results. Error bars represent the standard deviation of three technical triplicates.

**Figure S8.**
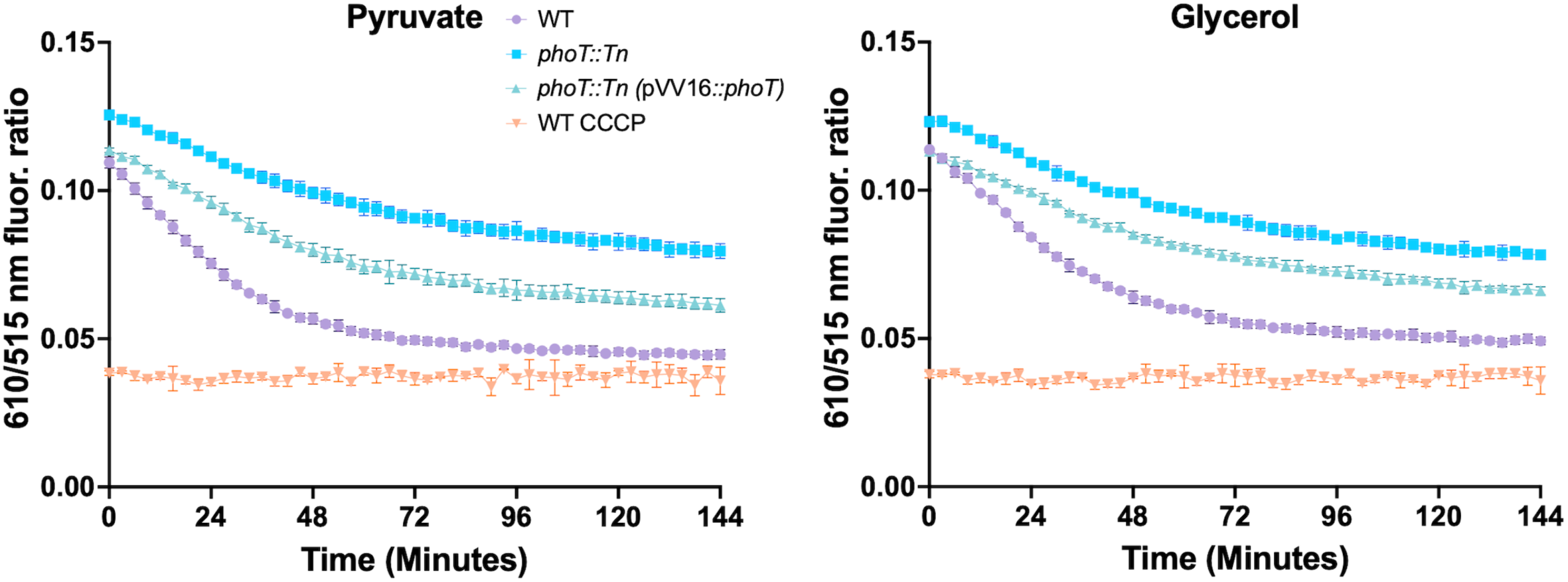
The *phoT::Tn* mutant has higher membrane potential in pyruvate and glycerol. Membrane potential of strains incubated in MMAT medium with high phosphate (25 mm) with different carbon sources. The strains were evaluated were *phoT::Tn* (pVV16::p*hoT*) (teal), *phoT::Tn* (light blue), and WT (purple). WT treated with CCCP (orange) was used as a depolarized control. This experiment was performed twice with similar results. Error bars represent the standard deviation of three technical replicates.

**Figure S9.**
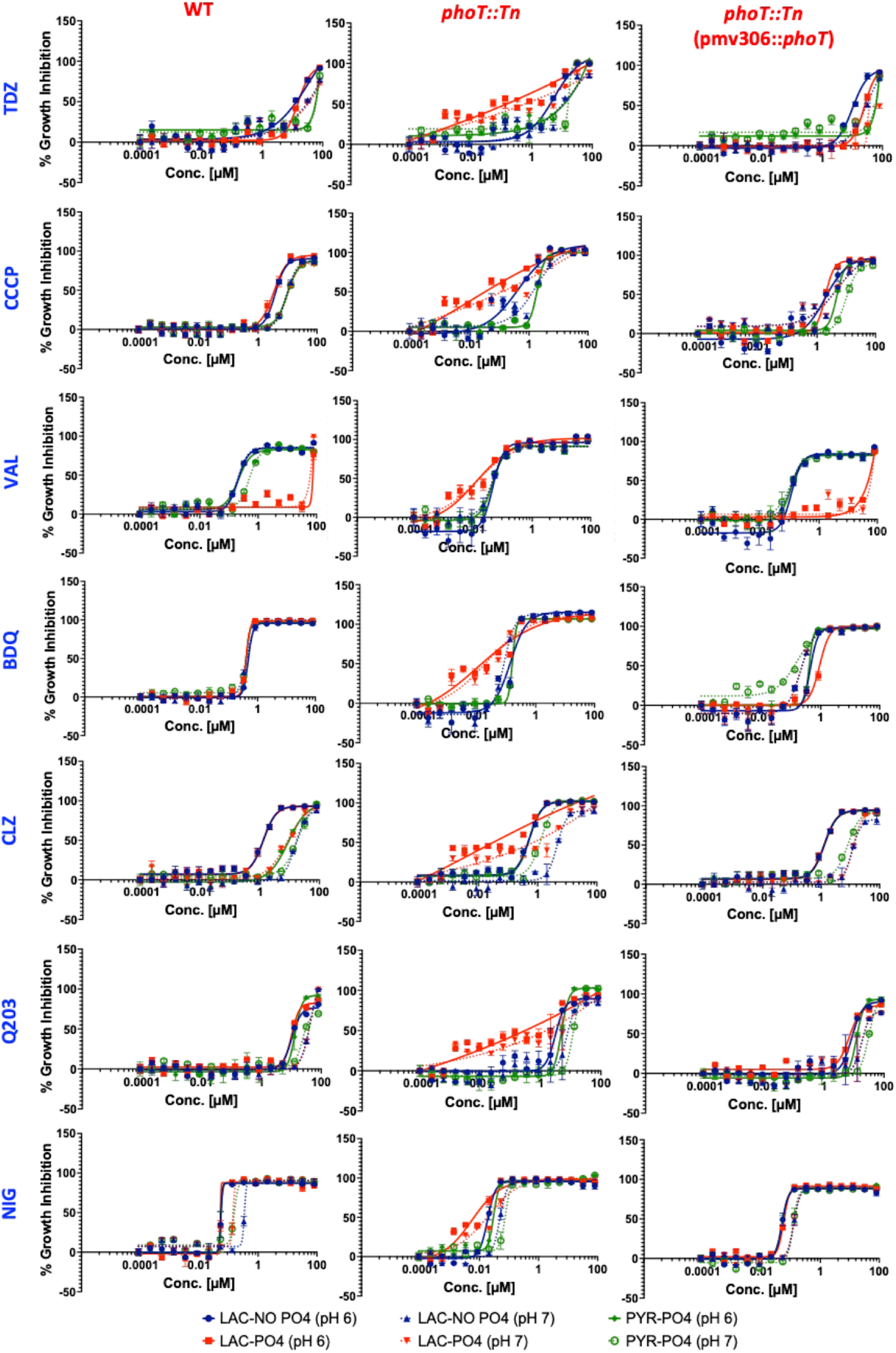
Dose-response curves of Electron transport chain-targeting agents across different growth conditions and genetic backgrounds. ETC-targeting agents were tested against WT, *phoT::Tn*, and *phoT::Tn* (pMV306::*phoT*) strains grown in different carbon sources (Lactate and Pyruvate), pHs (6 and 7), and in the presence and absence of phosphate to demonstrate the role of the lactate-induced ETC stress in potentiating the efficacy of ETC-targeting agents. TDZ: Thioridazine, CCCP: Carbonyl cyanide m-chlorophenyl hydrazone, VAL: Valinomycin, BDQ: Bedaquiline, CLZ: Clofazimine, NIG: Nigericin.

**Figure S10.**
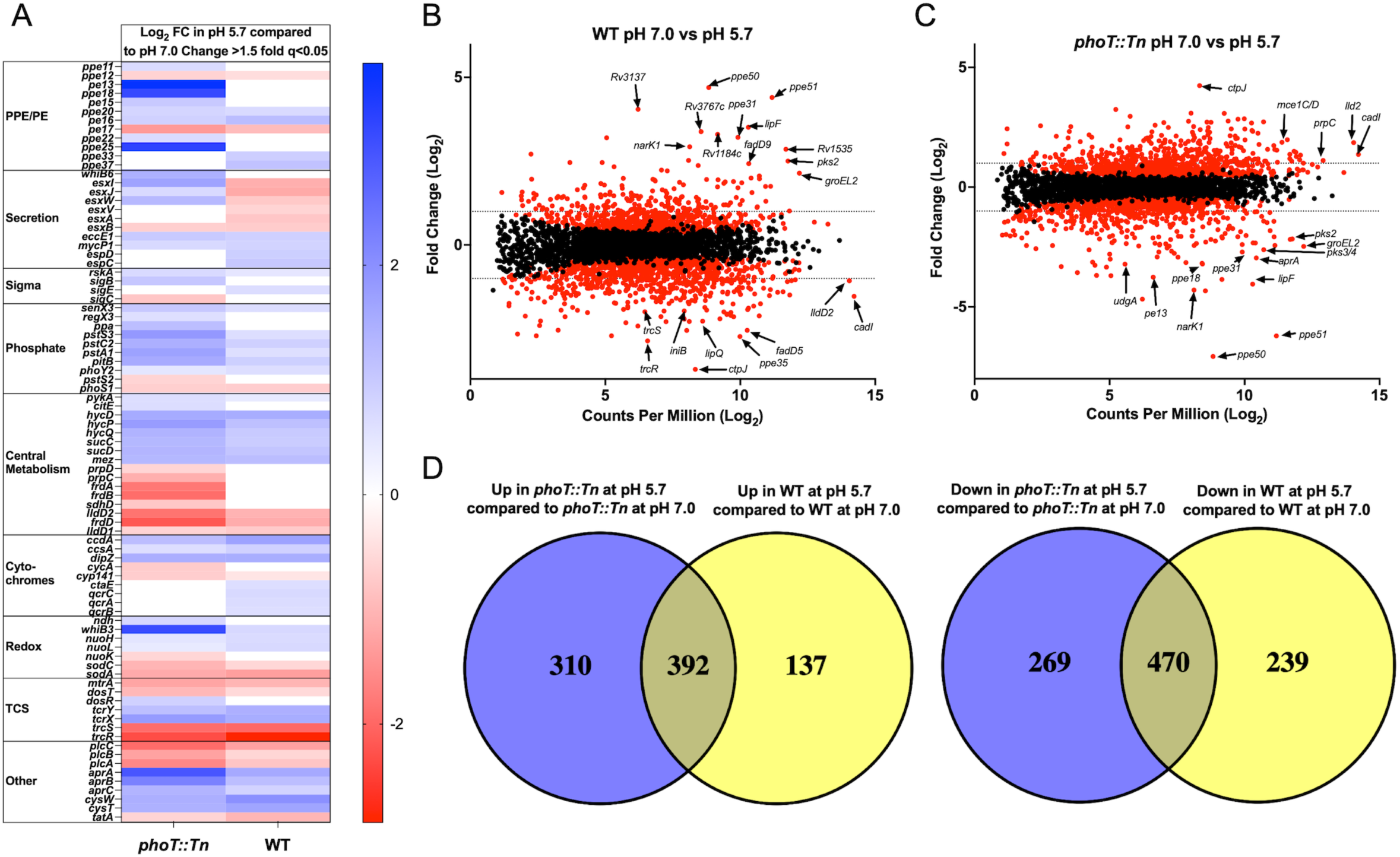
Genes differentially regulated when comparing pH 5.7 to 7.0 in the *phoT::*Tn mutant or WT Mtb. A) Heat map of genes differentially regulated (fold-change>1.5, q<0.05) in the *phoT::Tn* mutant at pH 5.7 compared to the *phoT::Tn* mutant at pH 7.0 (left) and WT grown at pH 5.7 compared to WT grown at pH 7.0 (right). B) Magnitude/amplitude plots of average Log_2_ counts per million (CPM) and Log_2_ fold-change of the *phoT::Tn* mutant grown at pH 5.7 compared to the *phoT::Tn* mutant grown at pH 7.0. C) Magnitude/amplitude plots of average Log_2_ counts per million (CPM) and Log_2_ fold-change of WT grown at pH 5.7 compared to WT grown at pH 7.0. D) Venn diagram depicting the overlap of genes upregulated in both strains at pH 5.7 compared to pH 7.0. Overlapping region indicates genes upregulated/downregulated in both strains at pH 5.7. Blue region indicates genes only upregulated in the *phoT::Tn* mutant at pH 5.7. The yellow region indicates genes only upregulated in WT at pH 5.7.

**Figure S11.**
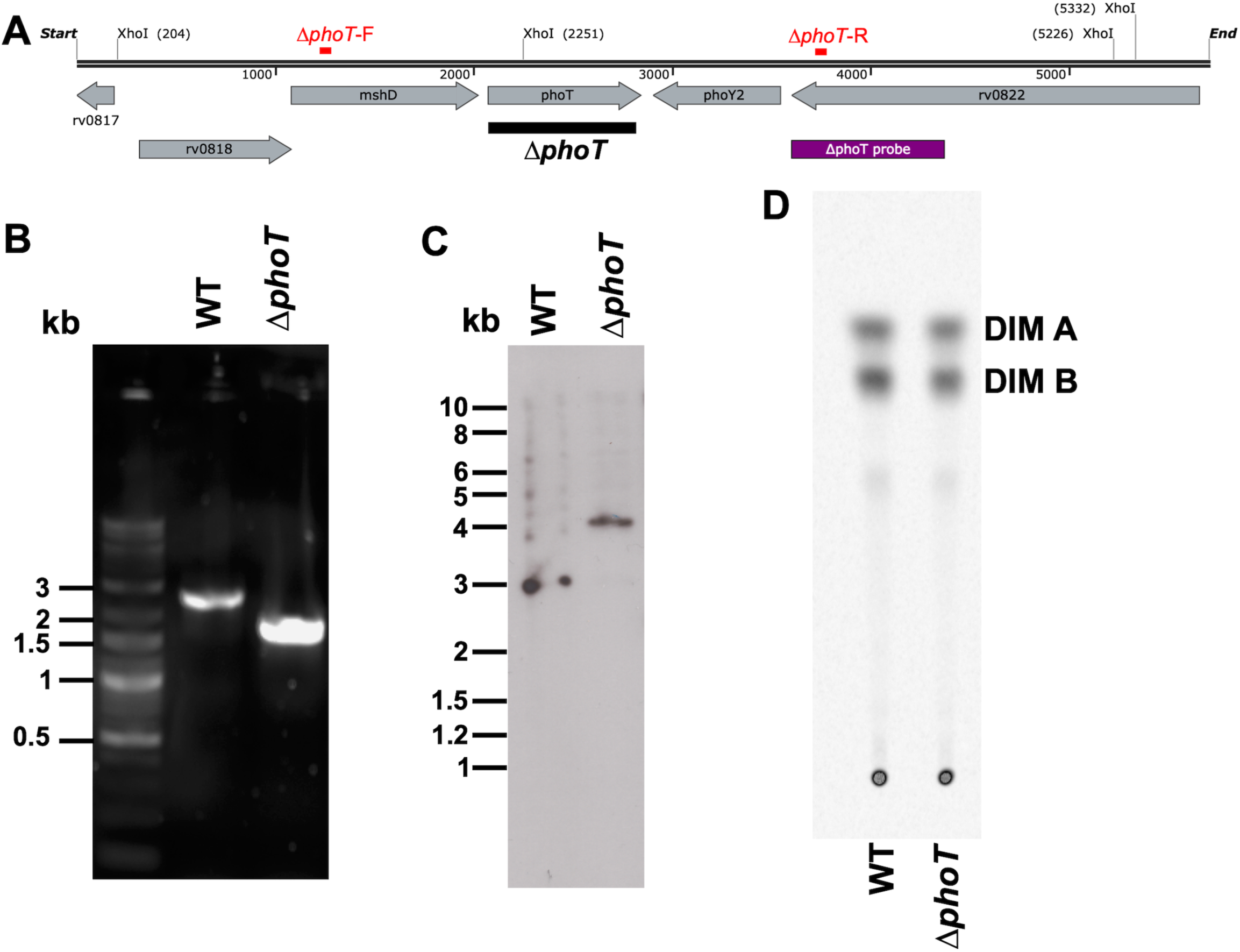
Δ*phoT* mutant validation. A) Diagram of the *phoT* locus. Black Δ*phoT* bar indicates the region deleted. Δ*phoT*-F and Δ*phoT*-R (red) are primers used to confirm the Δ*phoT* deletion by PCR. XhoI restriction enzyme sites and probe (purple) used for Southern blotting are also shown. B) PCR confirmation of the Δ*phoT* mutation using primers Δ*phoT*-F and Δ*phoT*-R on genomic DNA from WT and Δ*phoT*. Sizes of the DNA ladder in kb are indicated. C) Southern blotting confirmation of the Δ*phoT* mutation. Genomic DNA from WT and Δ*phoT* mutant *Mtb* was digested with XhoI, separated on a 0.8% TAE gel and transferred to a Hybond N+ nylon membrane (Amersham). A Δ*phoT* probe generated by PCR was labeled with the ECL direct nucleic acid labeling kit (Amersham). The blot was incubated with the probe overnight in Amersham Gold hybridization buffer with 0.5M NaCl and 5% blocking reagent at 42°C. Bands that hybridized to the probe were detected with ECL detection reagents (Amersham), autoradiographic film and an automated film processor. Positions of molecular size markers in kb are indicated. Δ*phoT* removes an XhoI site, so the expected size of the band is larger, ∼4.2 kb, as compared to ∼3.0 kb for WT. D) Phthiocerol dimycocerosate (PDIM) was detected in ^14^C propionate labeled apolar lipid extracts from WT and Δ*phoT* by thin layer chromatography. Spots corresponding to phthiocerol (methoxy; DIM A) and phthiodiolone (keto; DIM B) forms are indicated.

**Figure S12.**
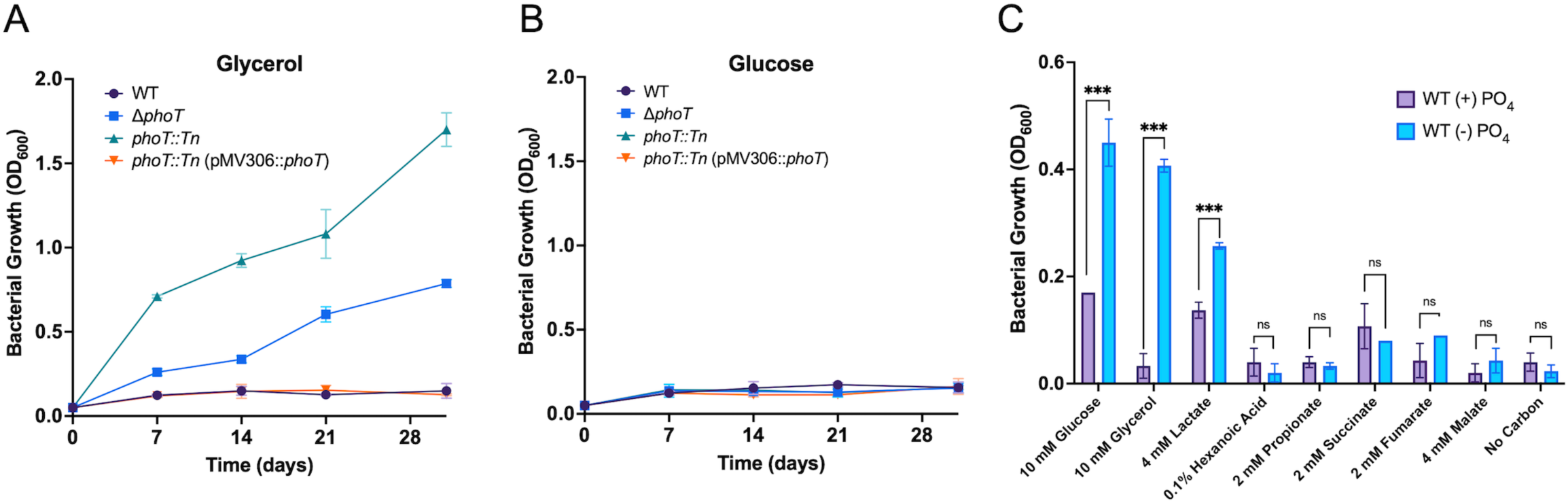
*phoT* mutants grow on glycerol, glucose, and lactate at acidic pH on modified 7H9 medium. A) *ΔphoT*, *phoT::Tn* mutant, WT and complemented strain growing at pH 5.0 on 0.2% (v/v) glycerol in carbon depleted 7H9. B) *ΔphoT*, *phoT::Tn* mutant, WT and complemented strain growth arrested on 2% (w/v) glucose at pH 5.0 in carbon depleted 7H9. C) Carbon panel of WT Erdman grown in MMAT at pH 5.7 in both phosphate rich medium, as well as phosphate replete medium. All three experiments were performed at least twice with similar results. Error bars represent the standard deviation of three technical replicates.

**Figure S13.**
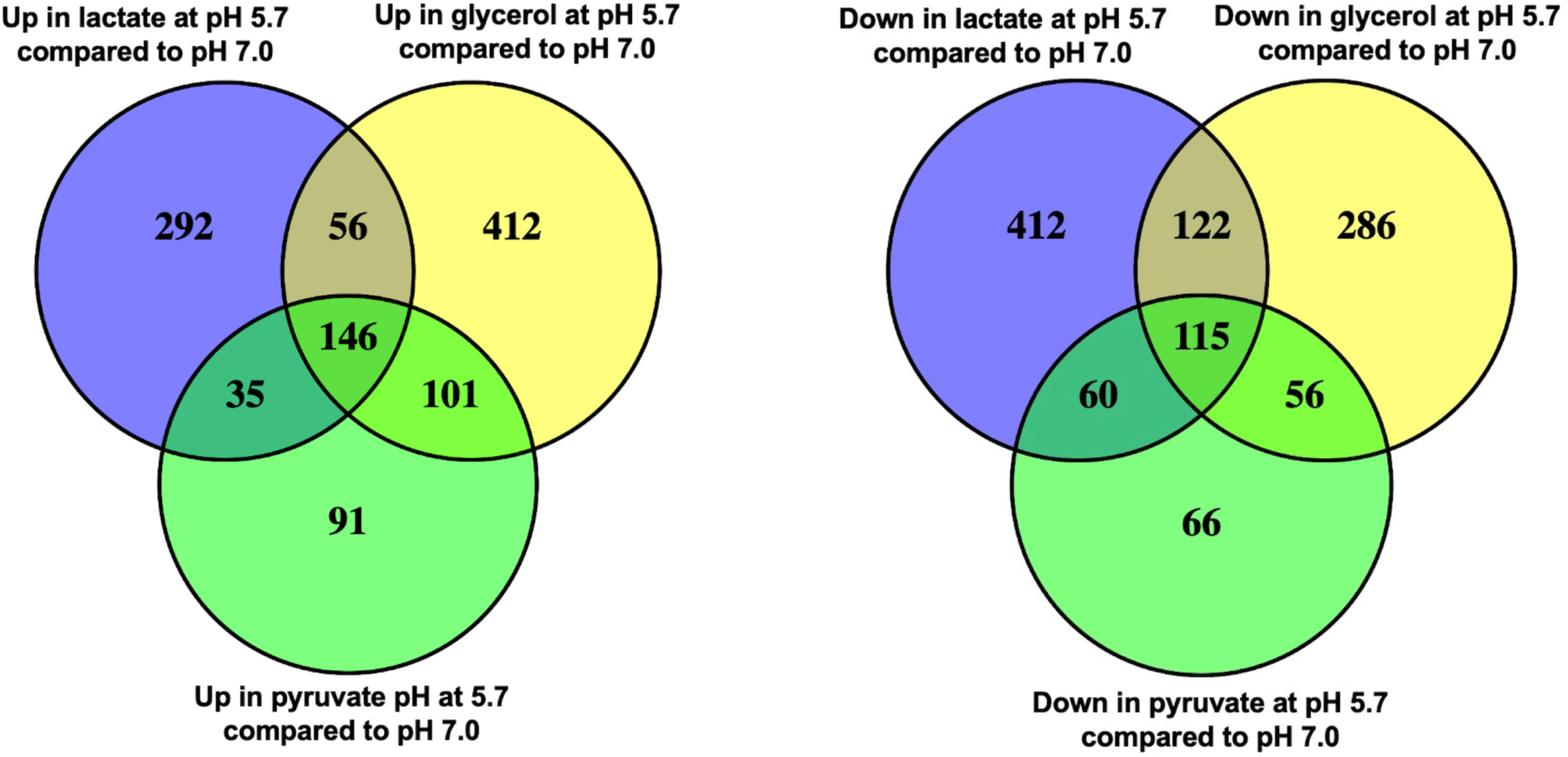
Subset of downregulated and upregulated genes are specific to lactate at acidic pH. Venn diagram depicting the overlay of genes upregulated in WT in either lactate, pyruvate, or glycerol at pH 5.7 compared to WT grown in the same carbon source at pH 7.0. The center depicts genes upregulated at acidic pH. The top left depicts genes upregulated only in lactate. The top right depicts genes only upregulated in glycerol. The bottom indicates genes only upregulated in pyruvate. In the venn diagram on the right, the center depicts genes downregulated at acidic pH in all conditions. The top left depicts genes downregulated only in lactate. The top right depicts genes only downregulated in glycerol. The bottom indicates genes only downregulated in pyruvate.

## References

1. Russell DG. 2011. Mycobacterium tuberculosis and the intimate discourse of a chronic infection. Immunol Rev 240:252–68.

2. Nadia L Ferrer ABG, Olivier Neyrolles, Brigitte Gicquel, Carlos Martin. 2010. Interactions of attenuated Mycobacterium tuberculosis phoP mutant with human macrophages. PLoS One 5.

3. Schaible UE, Sturgill-Koszycki S, Schlesinger PH, Russell DG. 1998. Cytokine activation leads to acidification and increases maturation o*f Mycobacterium avium*-containing phagosomes in murine macrophages. J Immunol 160:1290–6.

4. Martin C, Williams A, Hernandez-Pando R, Cardona PJ, Gormley E, Bordat Y, Soto CY, Clark SO, Hatch GJ, Aguilar D, Ausina V, Gicquel B. 2006. The live *Mycobacterium tuberculosis phoP* mutant strain is more attenuated than BCG and confers protective immunity against tuberculosis in mice and guinea pigs. Vaccine 24:3408–3419.

5. Vandal OH, Pierini LM, Schnappinger D, Nathan CF, Ehrt S. 2008. A membrane protein preserves intrabacterial pH in intraphagosomal *Mycobacterium tuberculosis*. Nat Med 14:849–854.

6. Abramovitch RB, Rohde KH, Hsu FF, Russell DG. 2011. *aprABC*: a *Mycobacterium tuberculosis* complex-specific locus that modulates pH-driven adaptation to the macrophage phagosome. Mol Microbiol 80:678–94.

7. Dechow SJ, Abramovitch RB. 2024. Targeting Mycobacterium tuberculosis pH-driven adaptation. Microbiology (Reading) 170.

8. Baker JJ, Johnson BK, Abramovitch RB. 2014. Slow growth of Mycobacterium tuberculosis at acidic pH is regulated by phoPR and host-associated carbon sources. Mol Microbiol 94:56–69.

9. Baker JJ, Dechow SJ, Abramovitch RB. 2019. Acid Fasting: Modulation of Mycobacterium tuberculosis Metabolism at Acidic pH. Trends Microbiol 27:942–953.

10. Baker JJ, Abramovitch RB. 2018. Genetic and metabolic regulation of Mycobacterium tuberculosis acid growth arrest. Sci Rep 8:4168.

11. Gouzy A, Healy C, Black KA, Rhee KY, Ehrt S. 2021. Growth of Mycobacterium tuberculosis at acidic pH depends on lipid assimilation and is accompanied by reduced GAPDH activity. Proc Natl Acad Sci U S A 118.

12. Dechow SJ, Baker JJ, Murto M, Abramovitch RB. 2022. ppe51 Variants Enable Growth of Mycobacterium tuberculosis at Acidic pH by Selectively Promoting Glycerol Uptake. J Bacteriol 204:e0021222.

13. Murdoch H, Dechow SJ, Abdalla BJ, Abramovitch RB. 2025. Mycobacterium tuberculosis growth arrest on propionate at acidic pH is suppressed by mutations in phoPR and pyrazinamide treatment. bioRxiv doi:10.1101/2025.09.24.678100.

14. Billig S, Schneefeld M, Huber C, Grassl GA, Eisenreich W, Bange FC. 2018. Lactate oxidation facilitates growth of Mycobacterium tuberculosis in human macrophages. Sci Rep 8:5241.

15. Jiang T, Gao C, Ma C, Xu P. 2014. Microbial lactate utilization: enzymes, pathogenesis, and regulation. Trends Microbiol 22:589–99.

16. Tischler AD, Leistikow RL, Kirksey MA, Voskuil MI, McKinney JD. 2013. Mycobacterium tuberculosis requires phosphate-responsive gene regulation to resist host immunity. Infect Immun 81:317–28.

17. Healy C, Ehrt S, Gouzy A. 2024. An exacerbated phosphate starvation response triggers Mycobacterium tuberculosis glycerol utilization at acidic pH. mBio doi:10.1128/mbio.02825-24:e0282524.

18. Ochman H, Gerber AS, Hartl DL. 1988. Genetic applications of an inverse polymerase chain reaction. Genetics 120:621–3.

19. Yunusbaeva M, Borodina L, Terentyeva D, Bogdanova A, Zakirova A, Bulatov S, Altinbaev R, Bilalov F, Yunusbayev B. 2024. Excess fermentation and lactic acidosis as detrimental functions of the gut microbes in treatment-naive TB patients. Front Cell Infect Microbiol 14:1331521.

20. Slonczewski JL, Fujisawa M, Dopson M, Krulwich TA. 2009. Cytoplasmic pH measurement and homeostasis in bacteria and archaea. Adv Microb Physiol 55:1–79, 317.

21. Sergine Even NDL, Pascal Loubière, Muriel Cocaign-Bousquet. 2002. Dynamic response of catabolic pathways to autoacidification in Lactococcus lactis: transcript profiling and stability in relation to metabolic and energetic constraints. Molecular Biology 45:1143–1152.

22. Alakomi HL, Skytta E, Saarela M, Mattila-Sandholm T, Latva-Kala K, Helander IM. 2000. Lactic acid permeabilizes gram-negative bacteria by disrupting the outer membrane. Appl Environ Microbiol 66:2001–5.

23. Amaral L, Kristiansen JE, Abebe LS, Millett W. 1996. Inhibition of the respiration of multi-drug resistant clinical isolates of Mycobacterium tuberculosis by thioridazine: potential use for initial therapy of freshly diagnosed tuberculosis. J Antimicrob Chemother 38:1049–53.

24. Lechartier B, Cole ST. 2015. Mode of Action of Clofazimine and Combination Therapy with Benzothiazinones against Mycobacterium tuberculosis. Antimicrob Agents Chemother 59:4457–63.

25. Stadler JAM, Maartens G, Meintjes G, Wasserman S. 2023. Clofazimine for the treatment of tuberculosis. Front Pharmacol 14:1100488.

26. Yano T, Kassovska-Bratinova S, Teh JS, Winkler J, Sullivan K, Isaacs A, Schechter NM, Rubin H. 2011. Reduction of clofazimine by mycobacterial type 2 NADH:quinone oxidoreductase: a pathway for the generation of bactericidal levels of reactive oxygen species. J Biol Chem 286:10276–87.

27. Lu P, Asseri AH, Kremer M, Maaskant J, Ummels R, Lill H, Bald D. 2018. The anti-mycobacterial activity of the cytochrome bcc inhibitor Q203 can be enhanced by small-molecule inhibition of cytochrome bd. Sci Rep 8:2625.

28. Pethe K, Bifani P, Jang J, Kang S, Park S, Ahn S, Jiricek J, Jung J, Jeon HK, Cechetto J, Christophe T, Lee H, Kempf M, Jackson M, Lenaerts AJ, Pham H, Jones V, Seo MJ, Kim YM, Seo M, Seo JJ, Park D, Ko Y, Choi I, Kim R, Kim SY, Lim S, Yim SA, Nam J, Kang H, Kwon H, Oh CT, Cho Y, Jang Y, Kim J, Chua A, Tan BH, Nanjundappa MB, Rao SP, Barnes WS, Wintjens R, Walker JR, Alonso S, Lee S, Kim J, Oh S, Oh T, Nehrbass U, Han SJ, No Z, et al. 2013. Discovery of Q203, a potent clinical candidate for the treatment of tuberculosis. Nat Med 19:1157–60.

29. Mahajan R. 2013. Bedaquiline: First FDA-approved tuberculosis drug in 40 years. Int J Appl Basic Med Res 3:1–2.

30. Andries K, Verhasselt P, Guillemont J, Gohlmann HW, Neefs JM, Winkler H, Van Gestel J, Timmerman P, Zhu M, Lee E, Williams P, de Chaffoy D, Huitric E, Hoffner S, Cambau E, Truffot-Pernot C, Lounis N, Jarlier V. 2005. A diarylquinoline drug active on the ATP synthase of Mycobacterium tuberculosis. Science 307:223–7.

31. Koul A, Dendouga N, Vergauwen K, Molenberghs B, Vranckx L, Willebrords R, Ristic Z, Lill H, Dorange I, Guillemont J, Bald D, Andries K. 2007. Diarylquinolines target subunit c of mycobacterial ATP synthase. Nat Chem Biol 3:323–4.

32. Wu Z, Bai L, Wang M, Shen Y. 2009. Structure–antibacterial relationship of nigericin derivatives. Chemistry of Natural Compounds 45:333–337.

33. Huang W, Briffotaux J, Wang X, Liu L, Hao P, Cimino M, Buchieri MV, Namouchi A, Ainsa JA, Gicquel B. 2017. Ionophore A23187 shows anti-tuberculosis activity and synergy with tebipenem. Tuberculosis (Edinb) 107:111–118.

34. Kevin Ii DA, Meujo DA, Hamann MT. 2009. Polyether ionophores: broad-spectrum and promising biologically active molecules for the control of drug-resistant bacteria and parasites. Expert Opin Drug Discov 4:109–46.

35. Chen S, Teng T, Zhang Z, Shang Y, Xiao H, Jiang G, Wang F, Jia J, Dong L, Zhao L, Chu N, Huang H. 2021. Carbonyl Cyanide 3-Chlorophenylhydrazone (CCCP) Exhibits Direct Antibacterial Activity Against Mycobacterium abscessus. Infect Drug Resist 14:1199–1208.

36. Cunarro J, Weiner MW. 1975. Mechanism of action of agents which uncouple oxidative phosphorylation: direct correlation between proton-carrying and respiratory-releasing properties using rat liver mitochondria. Biochim Biophys Acta 387:234–40.

37. Heytler PG. 1963. uncoupling of oxidative phosphorylation by carbonyl cyanide phenylhydrazones. I. Some characteristics of m-Cl-CCP action on mitochondria and chloroplasts. Biochemistry 2:357–61.

38. Andreoli TE, Tieffenberg M, Tosteson DC. 1967. The effect of valinomycin on the ionic permeability of thin lipid membranes. J Gen Physiol 50:2527–45.

39. Junge W, Schmid R. 1971. The mechanism of action of valinomycin on the thylakoid membrane : Characterization of the electric current density. J Membr Biol 4:179–92.

40. Su Z, Ran X, Leitch JJ, Schwan AL, Faragher R, Lipkowski J. 2019. How Valinomycin Ionophores Enter and Transport K(+) across Model Lipid Bilayer Membranes. Langmuir 35:16935–16943.

41. Parish T, Smith DA, Roberts G, Betts J, Stoker NG. 2003. The senX3-regX3 two-component regulatory system of Mycobacterium tuberculosis is required for virulence. Microbiology 149:1423–35.

42. Wren BW, Colby SM, Cubberley RR, Pallen MJ. 1992. Degenerate PCR primers for the amplification of fragments from genes encoding response regulators from a range of pathogenic bacteria. FEMS Microbiology Letters 99:287–291.

43. Himpens S, Locht C, Supply P. 2000. Molecular characterization of the mycobacterial SenX3-RegX3 two-component system: evidence for autoregulation. Microbiology (Reading) 146 Pt 12:3091–3098.

44. Glover R, Kriakov J., Garforth S, Baughn A, Jacobs W. 2007. The Two-Component Regulatory System senX3-regX3 Regulates Phosphate-Dependent Gene Expression in Mycobacterium smegmatis. Journal of Bacteriology 189:5495–5503.

45. Elliott SR, Tischler AD. 2016. Phosphate starvation: a novel signal that triggers ESX-5 secretion in Mycobacterium tuberculosis. Mol Microbiol 100:510–26.

46. White DW, Elliott SR, Odean E, Bemis LT, Tischler AD. 2018. Mycobacterium tuberculosis Pst/SenX3-RegX3 Regulates Membrane Vesicle Production Independently of ESX-5 Activity. mBio 9.

47. Ramakrishnan P, Aagesen AM, McKinney JD, Tischler AD. 2015. Mycobacterium tuberculosis Resists Stress by Regulating PE19 Expression. Infect Immun 84:735–46.

48. Wang Q, Boshoff HIM, Harrison JR, Ray PC, Green SR, Wyatt PG, Barry CE, 3rd. 2020. PE/PPE proteins mediate nutrient transport across the outer membrane of Mycobacterium tuberculosis. Science 367:1147–1151.

49. Mitra A, Speer A, Lin K, Ehrt S, Niederweis M. 2017. PPE Surface Proteins Are Required for Heme Utilization by Mycobacterium tuberculosis. MBio 8: e01720–16.

50. Sankey N, Merrick H, Singh P, Rogers J, Reddi A, Hartson SD, Mitra A. 2023. Role of the Mycobacterium tuberculosis ESX-4 Secretion System in Heme Iron Utilization and Pore Formation by PPE Proteins. mSphere 8:e0057322.

51. Ates LS, Ummels R, Commandeur S, van de Weerd R, Sparrius M, Weerdenburg E, Alber M, Kalscheuer R, Piersma SR, Abdallah AM, Abd El Ghany M, Abdel-Haleem AM, Pain A, Jimenez CR, Bitter W, Houben EN. 2015. Essential Role of the ESX-5 Secretion System in Outer Membrane Permeability of Pathogenic Mycobacteria. PLoS Genet 11:e1005190.

52. Kempker RR, Heinrichs MT, Nikolaishvili K, Sabulua I, Bablishvili N, Gogishvili S, Avaliani Z, Tukvadze N, Little B, Bernheim A, Read TD, Guarner J, Derendorf H, Peloquin CA, Blumberg HM, Vashakidze S. 2017. Lung Tissue Concentrations of Pyrazinamide among Patients with Drug-Resistant Pulmonary Tuberculosis. Antimicrob Agents Chemother 61:e00226–17.

53. B. S. Somashekarr DC. 2011. Metabolic Profiling of Lung Granuloma in Mycobacterium tuberculosis Infected Guinea Pigs: Ex vivo 1H Magic Angle Spinning NMR Studies. Journal of Proteome Research 10:4186–4195.

54. Stanley S, Wang X, Liu Q, Kwon YY, Frey AM, Hicks ND, Vickers AJ, Hui S, Fortune SM. 2023. Ongoing evolution of the Mycobacterium tuberculosis lactate dehydrogenase reveals the pleiotropic effects of bacterial adaption to host pressure. bioRxiv doi:10.1101/2023.10.09.561592.

55. Shi L, Salamon H, Eugenin EA, Pine R, Cooper A, Gennaro ML. 2015. Infection with Mycobacterium tuberculosis induces the Warburg effect in mouse lungs. Scientific Reports 5:18176.

56. Cumming BM, Pacl HT, Steyn AJC. 2020. Relevance of the Warburg Effect in Tuberculosis for Host-Directed Therapy. Frontiers in Cellular and Infection Microbiology 10.

57. Appelberg R, Moreira D, Barreira-Silva P, Borges M, Silva L, Dinis-Oliveira RJ, Resende M, Correia-Neves M, Jordan MB, Ferreira NC, Abrunhosa AJ, Silvestre R. 2015. The Warburg effect in mycobacterial granulomas is dependent on the recruitment and activation of macrophages by interferon-γ. Immunology 145:498–507.

58. Diep BA, Chan L, Tattevin P, Kajikawa O, Martin TR, Basuino L, Mai TT, Marbach H, Braughton KR, Whitney AR, Gardner DJ, Fan X, Tseng CW, Liu GY, Badiou C, Etienne J, Lina G, Matthay MA, DeLeo FR, Chambers HF. 2010. Polymorphonuclear leukocytes mediate Staphylococcus aureus Panton-Valentine leukocidin-induced lung inflammation and injury. Proc Natl Acad Sci U S A 107:5587–92.

59. Dinshaw KM, Lien KA, Knight M, Ouonkap SVY, Liu H, Savage DF, Carlson HK, Deutschbauer AM, Stanley SA. 2026. High-throughput characterization of Mycobacterium tuberculosis gene function across diverse conditions. PLoS Biol 24:e3003529.

60. Caslin HL, Abebayehu D, Pinette JA, Ryan JJ. 2021. Lactate Is a Metabolic Mediator That Shapes Immune Cell Fate and Function. Front Physiol 12:688485.

61. Jiang R, Ren WJ, Wang LY, Zhang W, Jiang ZH, Zhu GY. 2024. Targeting Lactate: An Emerging Strategy for Macrophage Regulation in Chronic Inflammation and Cancer. Biomolecules 14.

62. Kiran D, Basaraba RJ. 2021. Lactate Metabolism and Signaling in Tuberculosis and Cancer: A Comparative Review. Front Cell Infect Microbiol 11:624607.

63. Wang Y, Wu J, Lv M, Shao Z, Hungwe M, Wang J, Bai X, Xie J, Wang Y, Geng W. 2021. Metabolism Characteristics of Lactic Acid Bacteria and the Expanding Applications in Food Industry. Front Bioeng Biotechnol 9:612285.

64. Teusink B, Smid EJ. 2006. Modelling strategies for the industrial exploitation of lactic acid bacteria. Nat Rev Microbiol 4:46–56.

65. Mora-Villalobos JA, Montero-Zamora J, Barboza N, Rojas-Garbanzo C, Usaga J, Redondo-Solano M, Schroedter L, Olszewska-Widdrat A, López-Gómez JP. 2020. Multi-Product Lactic Acid Bacteria Fermentations: A Review. Fermentation 6.

66. Small JL, O’Donoghue AJ, Boritsch EC, Tsodikov OV, Knudsen GM, Vandal O, Craik CS, Ehrt S. 2013. Substrate specificity of MarP, a periplasmic protease required for resistance to acid and oxidative stress in Mycobacterium tuberculosis. J Biol Chem 288:12489–99.

67. Wilks JC, Slonczewski JL. 2007. pH of the cytoplasm and periplasm of Escherichia coli: Rapid measurement by green fluorescent protein fluorimetry. Journal of Bacteriology 189:5601–5607.

68. Martinez KA, 2nd, Kitko RD, Mershon JP, Adcox HE, Malek KA, Berkmen MB, Slonczewski JL. 2012. Cytoplasmic pH response to acid stress in individual cells of Escherichia coli and Bacillus subtilis observed by fluorescence ratio imaging microscopy. Appl Environ Microbiol 78:3706–14.

69. Cotter PD, Hill C. 2003. Surviving the acid test: responses of gram-positive bacteria to low pH. Microbiol Mol Biol Rev 67:429–53, table of contents.

70. Zhou C, Fey PD. 2020. The acid response network of Staphylococcus aureus. Curr Opin Microbiol 55:67–73.

71. Lund P, Tramonti A, De Biase D. 2014. Coping with low pH: molecular strategies in neutralophilic bacteria. FEMS Microbiol Rev 38:1091–125.

72. Weinrick B, Dunman PM, McAleese F, Murphy E, Projan SJ, Fang Y, Novick RP. 2004. Effect of mild acid on gene expression in Staphylococcus aureus. J Bacteriol 186:8407–23.

73. Richard H, Foster JW. 2004. Escherichia coli glutamate- and arginine-dependent acid resistance systems increase internal pH and reverse transmembrane potential. J Bacteriol 186:6032–41.

74. Rifat D, Bishai WR, Karakousis PC. 2009. Phosphate depletion: a novel trigger for Mycobacterium tuberculosis persistence. J Infect Dis 200:1126–35.

75. Mahatha AC, Mal S, Majumder D, Saha S, Ghosh A, Basu J, Kundu M. 2020. RegX3 Activates whiB3 Under Acid Stress and Subverts Lysosomal Trafficking of Mycobacterium tuberculosis in a WhiB3-Dependent Manner. Frontiers in Microbiology 11.

76. Larsson JT, Rogstam A, von Wachenfeldt C. 2005. Coordinated patterns of cytochrome bd and lactate dehydrogenase expression in Bacillus subtilis. Microbiology (Reading) 151:3323–3335.

77. Hsu T, Hingley-Wilson SM, Chen B, Chen M, Dai AZ, Morin PM, Marks CB, Padiyar J, Goulding C, Gingery M, Eisenberg D, Russell RG, Derrick SC, Collins FM, Morris SL, King CH, Jacobs WR, Jr. 2003. The primary mechanism of attenuation of bacillus Calmette-Guerin is a loss of secreted lytic function required for invasion of lung interstitial tissue. Proc Natl Acad Sci U S A 100:12420–5.

78. Piddington DL, Kashkouli A, Buchmeier NA. 2000. Growth of Mycobacterium tuberculosis in a defined medium is very restricted by acid pH and Mg(2+) levels. Infect Immun 68:4518–22.

79. Long JE, DeJesus M, Ward D, Baker RE, Ioerger T, Sassetti CM. 2015. Identifying essential genes in Mycobacterium tuberculosis by global phenotypic profiling. Methods Mol Biol 1279:79–95.

80. Sassetti CM, Boyd DH, Rubin EJ. 2001. Comprehensive identification of conditionally essential genes in mycobacteria. Proc Natl Acad Sci U S A 98:12712–7.

81. Kent WJ. 2002. BLAT—the BLAST-like alignment tool. Genome Res 12:656–64.

82. Deatherage DE, Barrick JE. 2014. Identification of mutations in laboratory-evolved microbes from next-generation sequencing data using breseq. Methods Mol Biol 1151:165–88.

83. Purdy GE, Niederweis M, Russell DG. 2009. Decreased outer membrane permeability protects mycobacteria from killing by ubiquitin-derived peptides. Mol Microbiol 73:844–57.

84. Rohde KH, Abramovitch RB, Russell DG. 2007. *Mycobacterium tuberculosis* invasion of macrophages: linking bacterial gene expression to environmental cues. Cell Host Microbe 2:352–64.

85. Johnson BK, Scholz MB, Teal TK, Abramovitch RB. 2016. SPARTA: Simple Program for Automated reference-based bacterial RNA-seq Transcriptome Analysis. BMC Bioinformatics 17:66.

